# CGGBP1-regulated cytosine methylation at CTCF-binding motifs resists stochasticity

**DOI:** 10.1101/2020.02.13.948604

**Authors:** Manthan Patel, Divyesh Patel, Subhamoy Datta, Umashankar Singh

## Abstract

The human CGGBP1 is implicated in a variety of cellular functions. It regulates genomic integrity, cell cycle, gene expression and cellular response to growth signals. Evidence suggests that these functions of CGGBP1 manifest through binding to GC-rich regions in the genome and regulation of interspersed repeats. Recent works show that CGGBP1 is needed for cytosine methylation homeostasis and genome-wide occupancy patterns of the epigenome regulator protein CTCF. It has remained unknown if cytosine methylation regulation and CTCF occupancy regulation by CGGBP1 are independent or interdependent processes. By sequencing immunoprecipitated methylated DNA, we have found that some transcription factor-binding sites resist stochastic changes in cytosine methylation. Of these, we have analyzed the CTCF-binding sites thoroughly and show that cytosine methylation regulation at CTCF-binding DNA sequence motifs by CGGBP1 is deterministic. These CTCF-binding sites are positioned at locations where the spread of cytosine methylation *in cis* depends on the levels of CGGBP1. Our findings suggest that CTCF occupancy and functions are determined by CGGBP1-regulated cytosine methylation patterns.

## INTRODUCTION

CTCF is chromatin architectural protein with a broad repertoire of functions [1]. It is regarded as a principal regulator of higher order chromatin structure. Maintenance of chromatin topology and chromatin boundaries are the key functions of CTCF [1–3]. The DNA-binding of CTCF is conventionally understood to take place through a consensus DNA sequence motifs. CTCF binds to different DNA sequence motifs like M1 and M2 [4], Ren_20 [5] and LM2, LM7 and LM23 [5,6]. Of the eleven Zn fingers in CTCF, one ZN7 interacts with cytosine in a methylation-sensitive manner [7]. This inhibition of CTCF-DNA binding and thus its function by motif methylation is a mechanism that regulates site-specific insulator activities of CTCF [8–12]. Methylation of CTCF-motifs and mitigation of CTCF function is a mechanism that has evolved to regulate the epigenome during development in a tissue-specific manner and has been reviewed extensively [13,14]. While a lot of research has addressed the functions and regulatory effects of CTCF, the regulation of CTCF by partnering proteins has remained less studied. CTCF-interacting proteins such as YY1, the Cohesin complex and BRD2 for example, cooperate with CTCF and are needed for its enhancer-promoter looping, topological domains maintenance and boundary element functions respectively [15–19]. However, what regulates methylation at CTCF-motifs remains largely unknown. The regulator of methylation at CTCF motifs would naturally also be a regulator of CTCF-DNA binding.

Recently we have demonstrated that the human CGG triplet repeat binding protein (CGGBP1) is required for normal genomic occupancy of CTCF [20]. CTCF not only binds to the DNA sequence specific motifs, but also to interspersed repeats, mainly L1-LINEs and Alu-SINEs [20]. In the presence of CGGBP1, the repeat occupancy of CTCF accounts for more than 40% of all the binding sites with L1 and Alu comprising the most of it. However, CGGBP1 depletion leads to an imbalance in the DNA-binding preference of CTCF. Upon CGGBP1 knockdown the repeat binding of CTCF is diminished and the CTCF-binding gets limited to the motifs [20]. Interestingly, like CTCF, CGGBP1 itself is a methylation-sensitive DNA-binding protein [20–25]. However, there is evidence that CGGBP1 binding to the target sequences prevents methylation from taking place [25,26]. WGBS experiments have shown that CGGBP1 depletion leads to genome-wide disturbances in methylation [27,28]. One of the major sites of methylation disturbances upon CGGBP1 knockdown are the binding sites of CGGBP1 itself, the Alu and L1 repeats [27,29]. There seems to be an evolutionary relationship between the CGGBP1-binding SINEs and CTCF binding sites [4] and the methylation regulation at Alu SINEs and CTCF-binding sites could thus have some hitherto unexplored evolutionary relationship as well. By maintaining the balance between CTCF at repeats or motifs, CGGBP1 acts as a regulator of CTCF binding pattern genome-wide. Data suggests that the repeat binding of CTCF takes place in cooperation with CGGBP1 [20]. Thus, while CGGBP1 depletion directly affects CTCF association with repetitive sequences, the gain of binding at motifs seems like an indirect consequence of CTCF displacement from the repeats. CGGBP1 however is also a methylation regulatory protein. Methylation changes at CTCF motifs can potentially affect CTCF binding and through it a change in the genome organization and function. We have previously shown that CGGBP1 depletion causes methylation disruption genome-wide with varied effects on repeats and sequence specific protein binding sites, including regions that contain enhancers and known as well as predicted CTCF-binding sites [28]. The current state of information about the regulation of CTCF binding by a methylation effector protein CGGBP1 suggests that CGGBP1-regulated methylation at CTCF-motifs could affect the binding of CTCF to motifs. The previous attempts to study the regulation of methylation by CGGBP1 using WGBS did not allow a high confidence detection of CTCF motifs in the sequence data [27,28]. These studies in CGGBP1-depleted human fibroblasts showed that although as a net change there was a mild increase in methylation, at most regions genome-wide there was a gain as well as a loss of methylation occurring at nearby cytosine residues.

Here we have used MeDIP-seq to analyze the cytosine methylation changes caused by CGGBP1 depletion in the same system of CT and KD HEK293T cells (in which we have recently demonstrated a regulation of CTCF occupancy by CGGBP1) as well as human foreskin fibroblasts (closely related to the 1064Sk cells in which we have previously studied CGGBP1-regulation of cytosine methylation by WGBS). We show that there is a widespread disruption of methylation caused by CGGBP1 depletion in both the cell types with stochasticity being a major feature of methylation patterns in CT and KD cells. We show that the stochasticity of methylation changes are partially explained by the allelic imbalances in methylation that occur upon CGGBP1 depletion. By a targeted analysis of transcription factor binding motifs in the JASPAR database, we report that CGGBP1 depletion disrupts methylation at a panel of transcription factors, including CTCF. We identify different cytosine methylation fates of CTCF-binding repeat-free motifs and motif-free repeats in CT and KD. Our analysis of MeDIP data in the flanks of RFMs show that these are CTCF binding sites required for maintenance of cytosine methylation patterns asymmetrically in the flanks of the CGGBP1-dependent CTCF-binding. These findings provide evidence that methylation regulation by CGGBP1 affects CTCF occupancy at RFMs and it is functionally relevant for cytosine methylation distribution *in cis.*

## RESULTS

### CGGBP1 depletion causes widespread stochastic changes in cytosine methylation

Non-targeting (CT) or CGGBP1-targeting (KD) shRNA were expressed in HEK293T cells as described elsewhere [20] (Fig S1). DNA fragments were enriched using methylcytosine antibody and the methylated DNA (hereafter the DNA with cytosine methylation is referred to as methylated DNA) sequenced on the Ion Torrent platform with mean read lengths of 150 bp. The alignment to hg38 was equally efficient in CT and KD (Table S1). Unlike WGBS described for control or CGGBP1-depleted cells earlier [27,28], the MeDIP captured only the methylated DNA and expectedly did not show any overall large differences in the sequence properties and base composition of CT and KD MeDIP DNAs (Table S2; show GC%, the percentage of CpG, CHG or CHH contexts and G/C skew). To characterize the differences in methylation between CT and KD, we compared the MeDIP signals (normalized read counts) for CT and KD in paired genomic bins (Fig 1A). The correlation between CT and KD MeDIP signals varied strongly with the genomic bin size for comparison of the MeDIP signals. At 10 Kb, the CT and KD methylation signals showed a near identity with a high Spearman correlation (Fig S2). However, with a progressive decrease in the bin size down to 0.2 Kb, the correlation was lost (Fig S2). Randomization of CT and KD reads showed that the correlation at higher bin size and a loss of correlation at lower bin sizes is not due to random differences in CT and KD MeDIP (Table S3). Difference upon sum ratios (Diff/Sum) were calculated for normalized methylation signals in genomic bins of 0.2 kb paired between CT and KD (Fig 1A). A frequency plot of the Diff/Sum values showed that there were large scale methylation disturbances genome-wide upon CGGBP1 depletion (Fig 1B). Interestingly, as reported before [28], strikingly similar magnitude and frequency of GoM and LoM were observed (Fig 1B).

**Figure 1.**
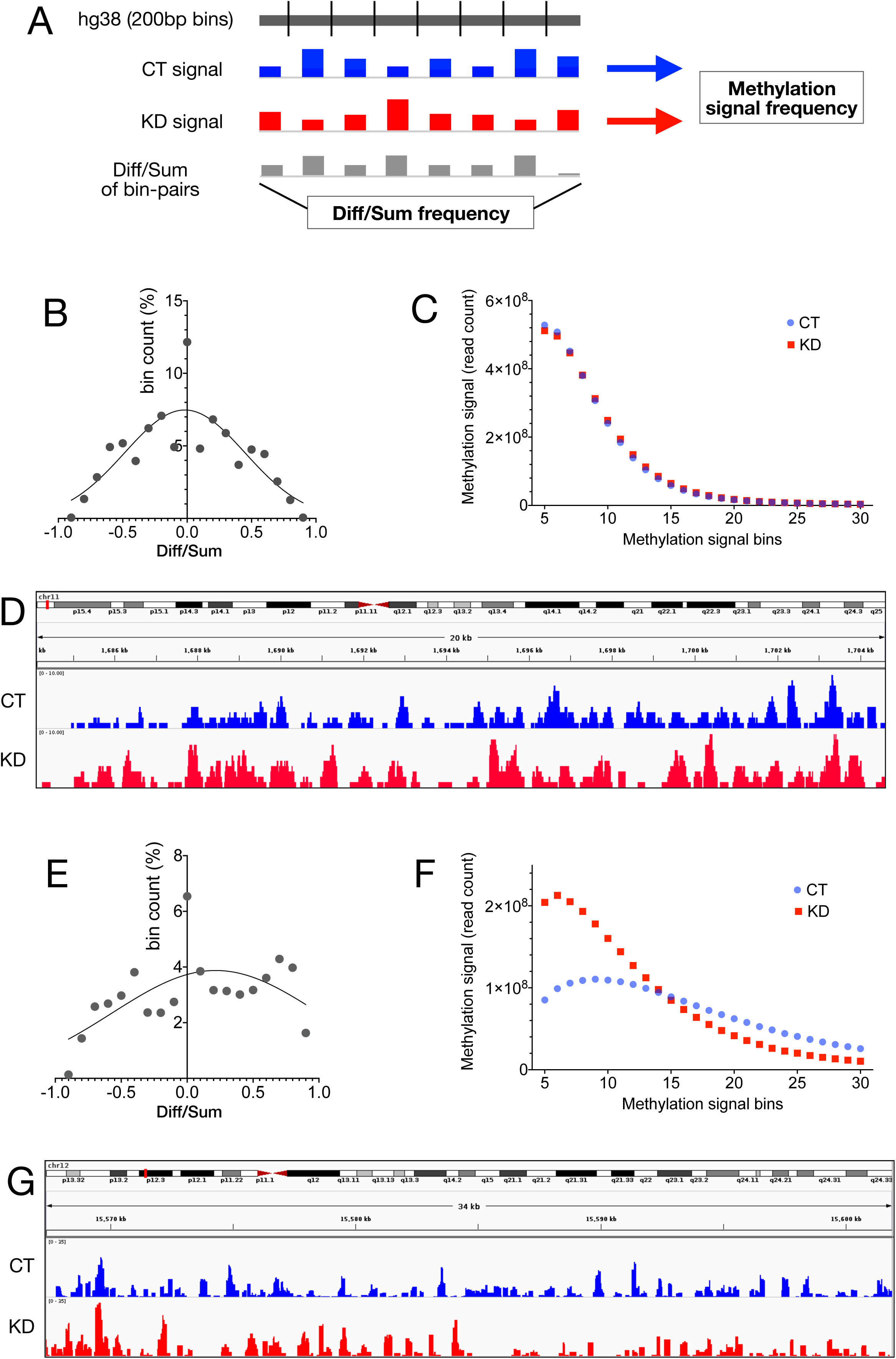
Stochastic cytosine methylation patterns are selectively dependent on CGGBP1 depletion. A: A schematic representation of the two main methods used to quantify and compare MeDIP signals from CT and KD. The normalized MeDIP signal was obtained for CT (blue) and KD (red) in 0.2 kb bins of hg38 and the Diff/Sum ([KD-CT]/[KD+CT]) ratios were calculated for each bin pair. The frequency plots of Diff/Sum ratios and MeDIP signals were used to compare CT and KD. B: Diff/Sum frequency in HEK293T shows a stochastic distribution resulting in a near-congruent gain-of-methylation (GoM) and loss-of-methylation (LoM). C: The widespread GoM and LoM (B) nullify any net change in cytosine methylation resulting in highly similar methylation frequencies in HEK293T CT and KD. D: Representative genome-view of HEK293T CT and KD MeDIP signals. E: The Diff/Sum ratio distribution for GM02639 has a skewed distribution showing a net GoM. F: The MeDIP signal distribution for GM02639 CT and KD show that the GoM and LoM are restricted to regions with low and high methylation levels respectively. G: A representative genome browser view of MeDIP signals for CT and KD.

For a quantitative assessment of the gross changes in methylation levels in HEK, as reported earlier in fibroblasts using WGBS, we binned the methylation signal genome-wide into discrete units of signals ranging from a minimum of 5 to a maximum of 30 (a genomic location represented at least 5 times to a maximum of 30 times respectively in the MeDIP-seq data) (Fig 1A). Consistent with the Diff/Sum distribution (Fig 1B), the methylation signal bin-wise distribution also revealed a near-identical distribution of methylation in CT and KD (Fig 1C). A representative genome browser view showed that both gain and loss of methylation indeed occurred at nearby locations (Fig 1D). We extended the MeDIP analyses to primary foreskin fibroblast GM02639 to relate the above mentioned findings in HEK293T with the previously reported methylation regulation by CGGBP1 in foreskin fibroblasts. Using siRNA against CGGBP1, we transiently knocked down CGGBP1 (Fig S3). In the previous studies, we have found the primary fibroblasts to be very sensitive to CGGBP1 depletion with a robust shutdown of transcription and exhibition of a stress-like phenotype [29]. To circumvent that, here we aimed at studying methylation changes caused by an incomplete depletion (approximately 50% knockdown by siRNA) of CGGBP1 in GM02639. We found that similar to HEK293T, the GM02639 also showed a widespread disturbance in methylation, however, with a net gain of methylation (Fig 1E). Similar to HEK, the correlation between CT and KD signals at 10 kb decreased drastically as we increased the methylation difference resolution to 0.2 kb (Fig S4). A frequency plot of the number of genomic regions represented for a range of methylation signals (from 5 to 30) showed that the representation of weakly methylated regions was strongly increased in KD (Fig 1F). This meant that the net increase in methylation was actually due to a much larger population of bins representing regions with low methylation signal in KD than in CT. A genome browser view of the representative positive as well as negative delta-signal regions showed that both gain and loss of methylation occurred at nearby locations (Fig 1G). These findings were similar to the previous methylation analyses in fibroblasts where CGGBP1 depletion showed coincidental gain and loss of methylation with a marginal net gain of methylation [27].

The contents of Satellite, Alu and L1 repeats were plotted against methylation changes as well as methylation signals. Unlike WGBS (which reports unmethylated as well as methylated cytosines), MeDIP (which captures and reports only what is methylated) expectedly pulled down repeats similarly in CT and KD. Interestingly, the LINE and Alu repeats showed mild but consistent methylation changes in HEK293T but only the satellite repeats were affected in GM02639 (Fig S5 and S6). Interestingly, although the GoM and LoM in HEK293T stochastically cancelled out each other, the methylation change analysis at LINE and Alu repeat subfamilies revealed specific changes. Some subfamilies exhibiting GoM and others undergoing LoM (Fig S5). The LINE and Alu repeats were affected in GM02639 only at very highly methylated regions (Fig S6). These included stochastic changes in methylation upon CGGBP1 depletion and methylation change at L1 repeats and their subfamilies.

We could not identify any sequence motifs or related sequence properties that were different between the CT or KD MeDIP DNA. We measured Shannon’s entropy for the probability distributions of any 0.2 Kb genomic bin (sequenced minimum 3 times in CT and KD combined and at least 2 times in either sample) to be different between CT and KD. For the range of differences which we used for Diff/Sum distributions, we plotted the entropies for the modulus of each Diff/Sum bin. For each of the bins the entropy values were calculated for the random probability of any region occurring in the state of no difference ((Diff/Sum) = 0) or difference ((Diff/Sum) = the bin value on X axis) between CT and KD. The entropy analyses showed that the overall entropy was very high for CT and KD of HEK293T as well as the GM02639 cells indicating a high stochasticity in methylation states. The stochasticity was however non-uniformly distributed for the GM02639 cells (Fig 2A). At higher magnitude of Diff/Sum ratios, the randomness was at its lowest suggesting that by applying a stringent cutoff for methylation change, we could extract the non-stochastic determinants of methylation change between CT and KD.

**Figure 2.**
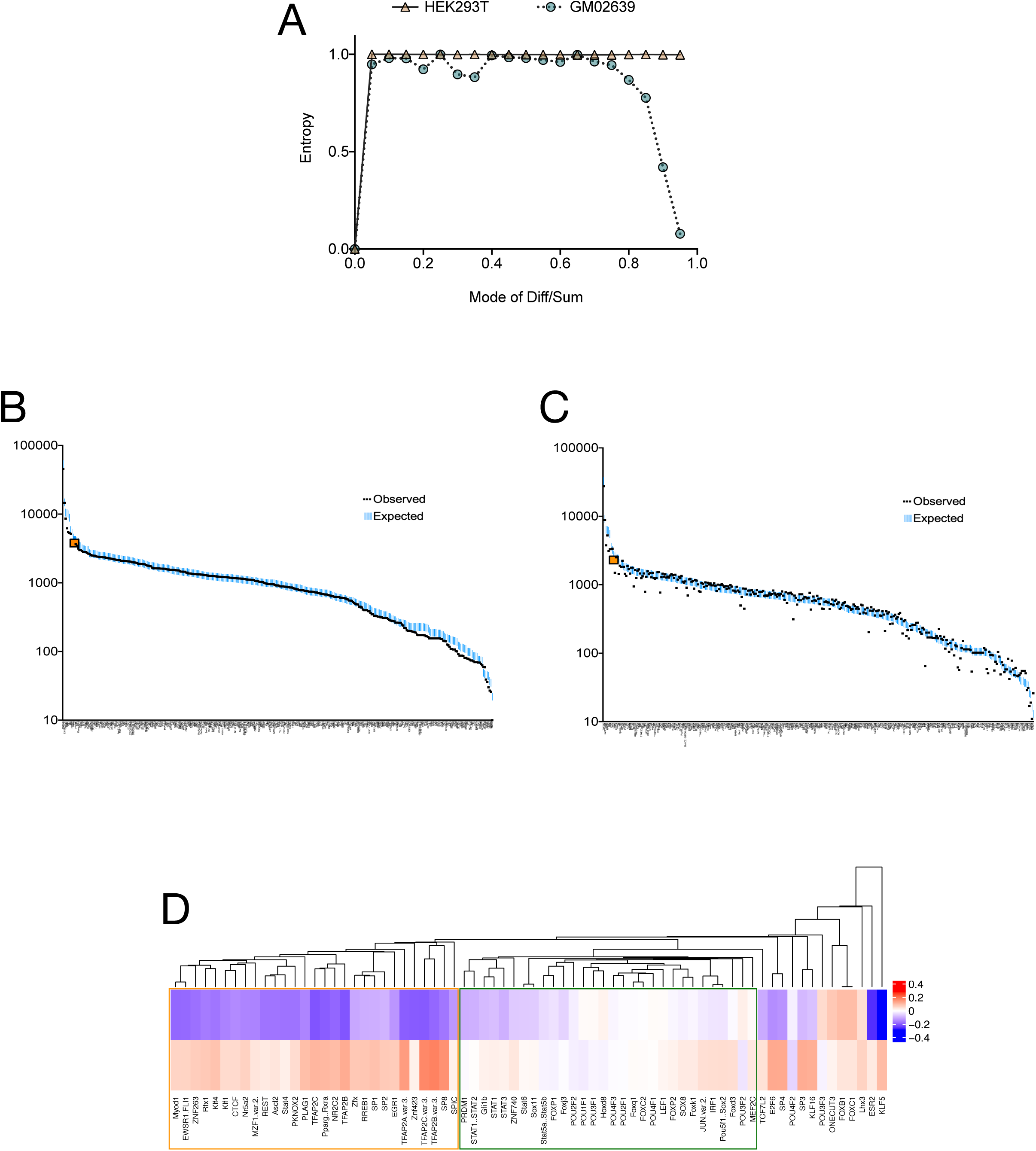
Methylation changes at specific transcription factor binding sites resist stochasticity. A: Shannon’s entropy distributions across the Diff/Sum bins show that the cytosine methylation changes in HEK293T and GM02639 have different levels of stochasticity. The HEK293T cells show a very high and uniform stochasticity for weak as well as strong methylation changes. In GM02639 however the stochastic methylation changes were weak. The strong changes in methylation were non-stochastic specifically in GM02639. This difference in stochasticity does not exclude the possibility that some genomic bins undergo methylation change commonly in HEK293T and GM02639. B: In the genomic bins sequenced (minimum sum of reads for CT and KD = 3) in CT and KD for HEK293T as well as GM02639 the JASPAR motifs occur with expected frequency showing that the MeDIP does not favour or exclude transcription factor binding sites (TFBS). C: In the bins undergoing net methylation change (|Diff/Sum| > 0.2) occurrence of the same JASPAR motifs as the ones called in B show deviations from the expected frequencies. The observed CTCF motif is highlighted in orange in B and C. D: A Chi^2^ test between the TFBS occurrences (B and C) identified a panel of 72 JASPAR motifs that are enriched in genomic bins differentially methylated between CT and KD. A single mode clustering classifies these motifs into two major groups: with opposite (orange box) or similar (green box) GoM and LoM between HEK293T and GM02639.

We thus applied a combination of three filters to extract and study deterministic changes in methylation: differentially methylated between CT and KD in HEK293T as well as GM02639, a minimum |Diff/Sum| value of 0.2 and a minimum sequence coverage of 3 reads per 0.2 Kb bin for CT and KD combined. Using these criteria we asked if the 0.2 Kb regions undergoing GoM or LoM are differentially enriched in DNA sequence motifs that constitute known protein binding sites.

### CGGBP1 regulated methylation patterns target multiple TFBSs including those of CTCF

Methylation inhibits DNA binding of most transcription factors (TFs) [30]. On the other hand, cytosine methylation favours binding of many proteins which do not have any DNA-sequence specificity [31]. We asked if the methylation disturbance caused by CGGBP1 depletion affects known transcription factor binding sites TFBSs.

A stringent search (p<1E-6) for JASPAR motifs of 519 TFs was performed in 0.2Kb bins covered in the CT or MeDIP dataset with a minimum coverage of 3 reads. This analysis showed that highly probable binding sites for more than 300 TFBSs are immunoprecipitated in MeDIP DNA of CT as well as KD (Fig 2B). In this search, the well known chromatin regulator protein CTCF featured as one of the proteins with highest occurrence in the CT and KD MeDIP DNA for both HEK293T and GM02639 (Fig 2B, orange data point) This constituted the expected frequency of TFBS occurrence in the combined MeDIP datasets. Subsequently, the TFBS frequencies were calculated in those genomic bins where the normalized methylation signals were different between CT and KD (|Diff/Sum| > 0.2) (Fig 2C, CTCF highlighted in orange). These observed TFBS frequencies for 343 TFs were compared against the expected frequencies and analyzed for each TF separately. A total of 72 TFs showed a significantly higher presence of TFBS in the observed (occurrence in bins differentially methylated between CT and KD) as compared to the expected. The methylation signal Diff/Sum ratios were calculated for these TFS separately in HEK293T and GM02639 datasets (Fig 2, B and C). Interestingly, most of these 72 TFs, including CTCF, showed opposite changes in methylation in HEK293T and GM02639 (Fig 2D).

CTCF binding to its motifs is regulated by cytosine methylation. CTCF occupancy at motifs as compared to repeats depends on the levels of CGGBP1 as has been demonstrated in HEK293T cells [20]. Whether the changes in cytosine methylation caused by CGGBP1 depletion play a role in determining CTCF binding to its motifs or its occupancy at repeats is not known. As a step towards understanding this possibility, we analyzed the methylation changes at CGGBP1-dependent CTCF-binding sites.

The nature of CTCF-binding DNA sequence motifs is different between CT and KD with a G/C weightage difference at position eight [20]. However, the CTCF motifs present in HEK293T GoM and LoM fractions (see Fig 1B) showed no such difference and resembled the canonical CTCF motif (Fig S7). The ChIP-seq reads which were pulled down in KD represented the motif-rich regions of the genome which remain bound to CTCF in the absence of CGGBP1 [20]. These regions, although motif-rich, are expected to maintain low methylation levels as compared to CT. We fetched these HEK293T CT and KD reads from the published CTCF ChIP-seq data [20] and measured methylation signals at them. As expected, the CTCF-bound CT and KD reads were distributed similarly in CT and KD MeDIP data with a concentration near low methylation bins (Fig 3A). These reads also gave rise to genuine peaks with CTCF-motifs, the distributions of which across the methylation bins also followed the same pattern as those of the reads (Fig 3B).

**Figure 3.**
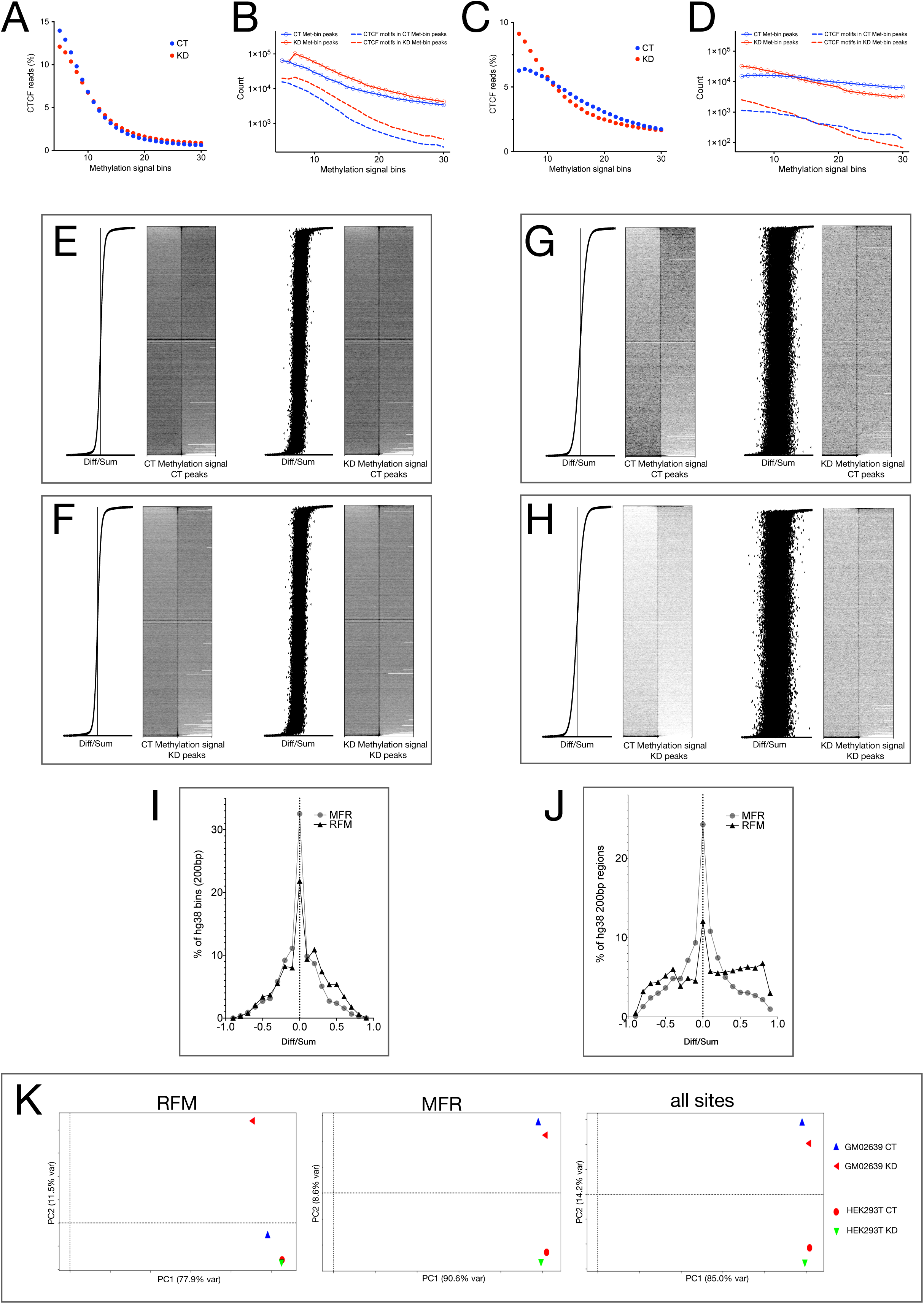
Cytosine methylation changes caused by CGGBP1 depletion are less stochastic at CTCF-binding RFM than at MFR. A: CTCF-binding sites in HEK293T are enriched in low methylation bins with no strong differences in MeDIP signals between CT and KD. B: MeDIP reads in HEK293T CT and KD are equally enriched in peaks (dashed lines) that are positive for CTCF motifs (continuous lines-circles). In agreement with the cytosine methylation-sensitivity of CTCF binding to DNA, the occurrence of peaks and CTCF motifs decline with increase in MeDIP signal. C: In GM02639 the concentration of CTCF-binding sites at low methylation bins is enhanced by CGGBP1 depletion. D: The occurrence of peaks in the GM02639 MeDIP reads (dashed lines) and their CTCF motif positivity (continuous lines-circles) regresses at a higher rate in KD showing that the cytosine methylation sensitivity of CTCF-binding at motifs in GM02639 is stronger in absence of CGGBP1. E and F: Cytosine methylation patterns at CTCF-binding sites in CT (E, n =) or KD (F, n =) are disrupted upon CGGBP1 depletion as revealed by MeDIP signals in the 10 kb flanks of the peak centres. G and H: GM02639 MeDIP signals at the same CTCF-binding sites (as shown in E and F) highlight three differences from the pattern observed in HEK293T (G and H compared with E and F respectively); overall low cytosine methylation levels, a stronger loss of methylation in KD compared to CT, and a larger disturbance in methylation caused by CGGBP1 depletion. A comparison of E and F with G and H respectively reinforces the findings that the cytosine methylation difference between CT and KD is less stochastic (Fig 2A) and the CTCF-binding motifs in KD are recused to a low methylation status in GM02639 (Fig 2, C and D, and Fig 3, C and D). I: The cytosine methylation changes observed in HEK293T (A, B, E and F) affect the repeat-derived (MFR) and motif-derived (RFM) CTCF-binding sites differently with a slight net GoM at RFM. J: The cytosine methylation changes in GM02639 CT and KD were strongly different between MFR (stochastic GoM and LoM with a normal distribution of Diff/Sum) and RFM (strong GoM and LoM with a multimodal distribution of Diff/Sum). A comparison of I and J shows that the cytosine methylation at RFM unlike MFR is specifically regulated by CGGBP1. The CGGBP1-dependence of RFM cytosine methylation is higher in GM02639 than HEK293T. K: PCA analyses reveal the different levels of stochasticity of methylation changes at RFM and MFR. The major component of variance (PC1) represented stochastic changes as it failed to segregate the CT and KD samples of the two cell types when RFM and MFR were analyzed separately or together. Commensurate with the previously described findings, the PC1 accounted for the least stochastic variance in RFM (77.9%) and highest (90.6%) for MFR. The PC2 accounted for 11.2% of variance between GM02639 CT and KD RFM only. For MFR and all sites, the PC2 accounted for variances between the cell types but not CT versus KD. Thus, at RFM the MeDIP signals have a GM02639-specific dependence on CGGBP1 whereas the same at MFR follow a cell type-specific pattern predominantly.

The findings with the JASPAR-wide motif search (Fig 2C) showed that the effect of CGGBP1 depletion on CTCF motifs in GM02639 would be different from that observed in HEK293T. The MeDIP signals were then used to identify how the methylation patterns were affected in GM02639 at the CTCF-bound DNA in CT and KD. Indeed the reads with low methylation signals were highly increased in KD. Strikingly, the identification of genuine peaks and motifs in those peaks was also restricted to low methylation bins in KD as compared to CT (Fig 3C and D). These results showed that motifs containing CTCF-binding sites are sensitive to CGGBP1-dependent methylation in HEK293T and GM02639 to varying degrees.

A panel of demonstrated and possible CTCF-binding sites were selected for analyzing methylation changes caused by CTCF depletion to validate the finding that methylation at CTCF-binding sites are regulated by CGGBP1. By using qPCR on fibroblast CT and KD genomic DNA digested with methylation-sensitive or methylation dependent restriction endonucleases, we established that methylation levels at many CTCF-binding sites are indeed affected by CGGBP1 depletion (Fig S8). We then cross-validated these findings by analyzing the CT and KD methylation signals at the previously characterized CGGBP1-regulated CTCF-binding sites. The disturbances observed in methylation patterns at CT peaks (Fig 3E) or KD peaks (Fig 3F) for HEK293T were weaker than the same observed for GM02639 CT (Fig 3G) or KD peaks (Fig 3 H). Noticeably, in GM02639, where methylation at CTCF-binding sites and flanks were strongly affected by CGGBP1 depletion, the binding of CTCF was restricted in KD to regions with much lower methylation levels than CT (Fig 3G compared with Fig 3H). Overall, the occurrence of CTCF-binding sites in GoM or LoM regions as well as occurrence of GoM and LoM at CTCF-binding sites together established that CGGBP1 depletion causes targeted methylation changes at CTCF-binding sites and its flanks in a cell type specific manner. However, CTCF-binding sites at repeats and motifs show opposite changes in CTCF occupancy upon CGGBP1 depletion. If methylation regulation by CGGBP1 is a potential means to regulate CTCF binding, then CGGBP1 depletion would cause different patterns of methylation changes at CTCF-binding repeats and motifs. We thus performed a comparative analysis of methylation change at CTCF-binding repeats and motifs.

### CGGBP1 affects methylation at CTCF-binding repeat-free motifs and motif-free repeats differently

We have described that the motif-free repeats (MFR) and repeat-free motifs (RFM) constitute exclusive sets CGGBP1-regulated CTCF-binding sites [20]. Whereas CTCF occupancy at MFR depends on CGGBP1, the same at RFM is not clearly understood. As described above, methylation changes caused by CGGBP1 depletion at CTCF-binding sites are concentrated at motifs with no specific changes at the repeats. To test this more rigorously, we focussed on MFR and RFM for a comparative analysis of methylation changes caused by CGGBP1 depletion at CTCF-binding sites. The methylation signals at RFM and MFR were compared and Diff/Sum values calculated for paired bins between CT and KD. Methylation disturbances were normally distributed in HEK293T at MFR as well as RFM. Unlike MFR, the M value distribution in RFM showed a slight positive skew (Fig 3I), which was in agreement with the findings of methylation change at CTCF motifs described in figure 3A. In GM02639, the Diff/Sum distribution of methylation changes at MFR were normally distributed with an approximately 30% reduction in the count of bins showing no methylation change (Fig 3J) compared to that in HEK293T (Fig 3I). The fraction of RFM undergoing a methylation change with |Diff/Sum| = 0.5 was more than two folds higher in GM02639 than in HEK293T. The M value distribution of RFM in GM02639 however showed a clear deviation from normal distribution with three modes and with a marked increase in the non-zero Diff/Sum population, which represents the fraction undergoing methylation change (Fig 3J). These results were also consistent with the findings that methylation changes in GM02639 due to CGGBP1 depletion were more pronounced than the same in HEK293T.

The Diff/Sum distributions of stochastic changes in methylation are expected to be normal. The deviations from a normal distribution indicate a specific association between RFM and methylation change in KD as compared to CT. The net methylation changes are however a sum of stochastic changes and specific changes. We performed PCA analysis to find out how the RFM and MFR methylation changes in CT and KD are different between HEK293T and GM02639.

As shown in figure 3K, the largest principal component that accounted for most of the variance failed to differentiate either the two cell lines or the samples CT and KD. The percentage of variance accounted for by this major component was 78% at RFM and 90% at MFR and 85% when all methylation changes across the genome (hg38) were measured (Fig 3K). This large component of variance not accounting for differences between the samples reflected the stochasticity of methylation changes. The second principal component accounted for the variation between GM02639 CT and KD at RFM only. For MFR and hg38, the second principal component only accounted for differences between the two cell lines. Thus, the difference between the methylation patterns at CT RFM and KD RFM in GM02639 was the only non-stochastic change in methylation across all the combinations of RFM, MFR and CT or KD in the two cell lines. Down to the fourth principal component (accounting for > 99.99% of variance) the CT and KD could not be differentiated at MFR in either cell line (not shown).

These analyses confirmed that CGGBP1 regulates methylation at RFM in GM02639 specifically. We argued that a specific regulation of methylation at RFM in GM02639 should also manifest as a non-stochastic and predictable pattern of methylation change between CT and KD at RFM and not MFR. To pursue this, we compared the methylation patterns in the flanking regions of the RFM and MRF.

### CTCF-binding RFM correspond to methylation transition boundaries

CTCF binding at the MFR has been shown to be required for restriction of H3K9me3 spread. Ablation of CTCF binding at repeats results in a disruption in H3K9me3 levels in the flanks of the MFR with most MFR exhibiting a loss of barrier activity (LoB) upon CGGBP1 depletion. The current findings that unlike MFR, the RFM are specifically associated with cytosine methylation changes suggested that similar to the barrier activities of MFR for H3K9me3 levels, the RFM could act like barriers for cytosine methylation levels. The difference between upstream and downstream methylation signals in 10 kb flanks of RFM and MFR were calculated for CT and KD HEK293T. There was a widespread asymmetry in the methylation signals obtained from upstream and downstream flanks of RFM (Fig 4A). The methylation level asymmetries in RFM flanks were poorly correlated between CT and KD (Fig 4B). On the other hand, the same analysis for MFR showed that the asymmetries between methylation signals in the upstream and downstream flanks of MFR were higher than those observed for RFM (Fig 4C; compared with Fig 4A), yet highly correlated between CT and KD. These findings in MFR were commensurate with a more stochastic change in methylation in MFR flanks as compared to RFM. Visualization of methylation signals in HEK293T (Fig 4, E to H) showed that at some RFM a lower downstream (Fig 4E) or upstream (Fig 4F) methylation in CT is lost in KD. Similarly, a gain of methylation selectively upstream (Fig 4G) or downstream (Fig 4H) of CTCF-binding sites was observed at other MFR. This methylation level asymmetry in the flanks of RFM was expected to be more widespread in GM02639. Indeed a visualization of methylation signals in GM02639 CT and KD showed that a larger number of RFM showed a loss of methylation asymmetry due to an increase of methylation in the downstream (Fig 4I) or upstream (Fig 4J) flanks in KD. An even larger number of RFM showed a gain of methylation asymmetry due to an increase in methylation selectively in the upstream (Fig 4K) or downstream (Fig 4L) flanks in KD. These predictable and deterministic methylation changes occurring selectively at RFM could be of functional consequence for CTCF binding to motifs and regulation of chromatin barrier activities. Interestingly, the RFM regions with cytosine methylation asymmetries were different from the repeat-rich CTCF binding sites that act as barriers for H3K9me3 levels [20] and we could not find any overlap between them (not shown).

**Figure 4:**
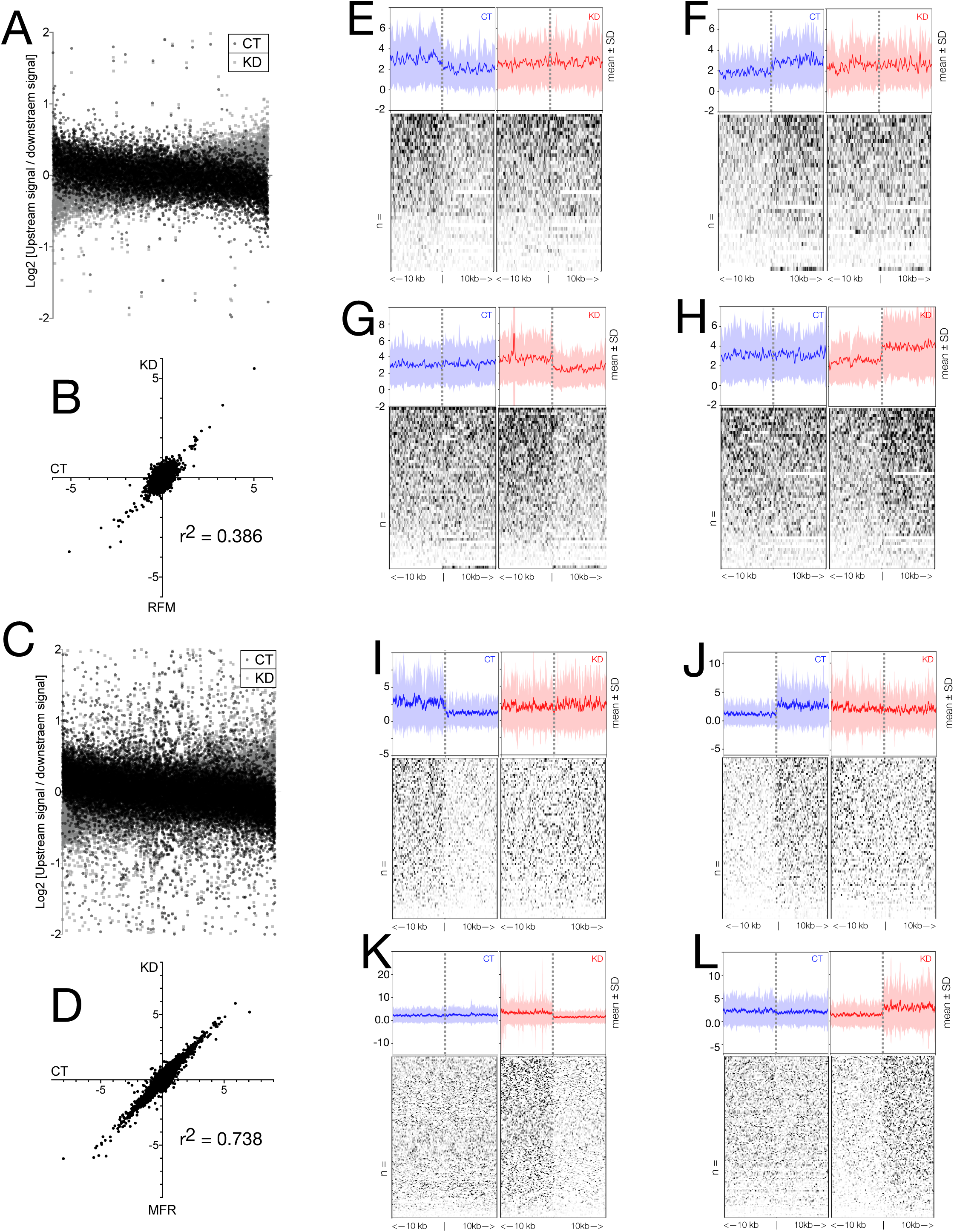
CGGBP1-dependent regulation of cytosine methylation spread by RFM CTCF-binding sites. A to D: Cytosine methylation signals in 10 kb flanks of RFM or MFR CTCF-binding sites suggest a barrier-like function for RFM. Upstream and downstream MeDIP-signals at RFM (A) show a stronger asymmetry (Pearson r2 = 0.386) (B) than that at MFR (C), which are better correlated (Pearson r2 = 0.738) and symmetrical (D). E to H: Cytosine signal asymmetry at HEK293T RFM is lost due to an increase in cytosine methylation downstream (E) or upstream (F) of the CTCF-binding sites. Conversely, the asymmetry is gained due to a decrease in cytosine methylation downstream (G) or upstream (H). I to L: The cytosine methylation asymmetry in RFM flanks in GM02639 is stronger than that observed in HEK293T and is lost due to an increase in methylation downstream (I) or (J). Similarly, a much stronger cytosine methylation asymmetry is achieved in GM02639 RFM than the HEK293T RFM due to a decrease in methylation downstream (K) or upstream (L). The stronger asymmetry of MeDIP signal in RFM flanks of GM02639 as compared to HEK293T reinforce a lower stochasticity of methylation change caused by CGGBP1 depletion in the former.

These sites for specific methylation regulation by CGGBP1 however were embedded in a much larger fraction of the genomic regions at which methylation changes were stochastic. One possibility that could explain this stochastic methylation disruption is that upon CGGBP1 depletion the two allelic copies become amenable to methylation changes independently. Allele-specificity of CTCF-binding and its regulation by allele-specific methylation is an established mechanism. We wanted to find out if unexpected levels of allelic imbalance underlies the stochasticity in methylation changes caused by CGGBP1 depletion. Thus, we analysed the allelic imbalance in methylation and its occurrence in CT and KD MeDIP datasets.

### Unexpected levels of allelic imbalance in MeDIP DNA upon CGGBP1 depletion

To find out the contribution of allelic imbalances towards stochasticity of methylation changes observed upon CGGBP1 depletion, we studied the proportions of alleles represented in MeDIP data separately for HEK293T and GM02639.

Allelic counts were obtained in HEK293T input [20], and compared with MeDIP CT and KD for all loci where both or either samples are heterozygous. The homozygosity or heterozygosity was called only if the locus was covered minimum five times in each sample. The presence of reference (Ref) and alternate (Alt) alleles constituted a heterozygous genotype, whereas occurrence of only Ref or Alt was called as homozygous. If CGGBP1 depletion did not affect methylation with an allelic bias, then the Alt and Ref genotypes would be represented equally in CT and KD and the overall genotype distributions for CT and KD MeDIP DNA would resemble that of the input. The input for HEK293T showed an expected skewed distribution with a higher presence of the Ref allele as compared to the Alt allele. The CT and KD MeDIP however showed a multimodal distribution with an unexpectedly high representation of the Ref as well as the Alt alleles (Fig 5A). The observed allelic distributions in CT and KD were both disturbed and different from the expected distribution of alleles as seen in the input (Fig 5, B and C), but were highly similar to each other (Fig 5D). Thus, the CT and KD MeDIP DNA had an unexpected genotype distribution clearly demonstrating an allelic imbalance. Also, the near congruence of the allelic ratio distributions showed that the deviation from the expected genotype ratios was stochastic.

**Figure 5:**
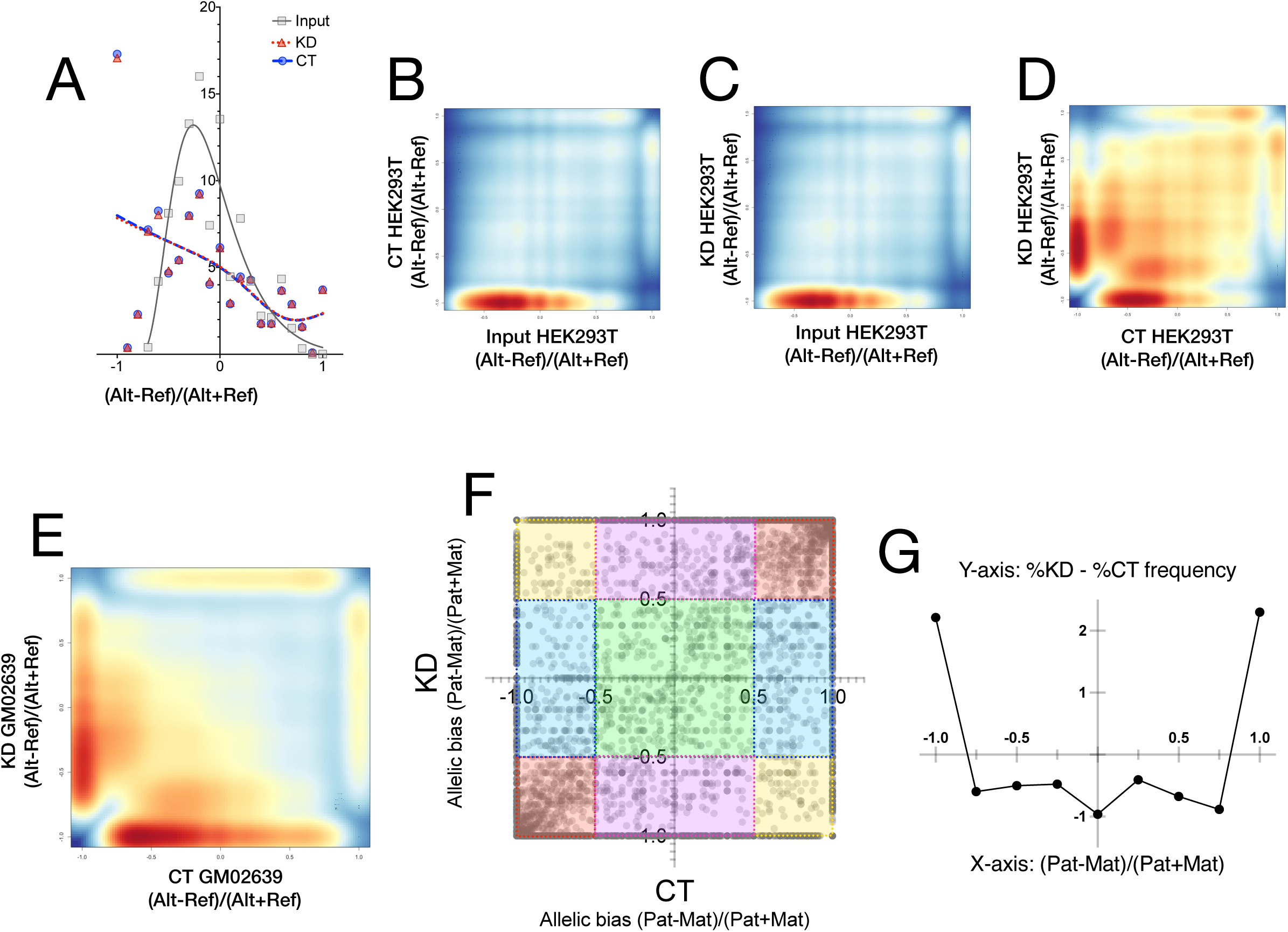
Stochastic allelic imbalances in cytosine methylation are exacerbated by CGGBP1 depletion. A: Allelic imbalances in HEK293T MeDIP are strongly different from the allelic ratios expected in the input with negligible differences between CT and KD. The poor correlation between allelic imbalances in the input and CT (B) or KD (C) is evident in the heat scatters (r2 = for B and for C). The stochastic allelic choices for methylation give rise to a random distribution of allelic representations in HEK293T CT and KD MeDIP (D). The allelic imbalances between GM02639 CT and KD were however highly anticorrelated as expected due to less stochasticity in these samples. F and G: Parent-of-origin specific allelic imbalance scatter plot between GM02639 CT and KD (F) shows that CGGBP1 depletion enhances a random monoallelic bias with no preference for the maternal or the paternal alleles. For all locations where the CT and KD are expected to be heterozygous ([(Pat-Mat)/(Pat+Mat)] values = 0±0.5), the observed heterozygosity was very low (green shade; see text for details). Instead the unbiased monoallelic methylation with no difference between CT and KD ([(Pat-Mat)/(Pat+Mat)] values < −0.5 or > 0.5) were observed for the majority of loci (red shades). A skewed allelic imbalance only in CT (aqua shade) or KD (pink shade). For a proportionate subset of loci, the parent-of-origin identities of the methylated alleles were reversed between CT and KD (yellow shades). G: A frequency plot of the differences between percent of maternal and paternal allele contributions to the MeDIP DNA shows that CGGBP1 depletion minimizes allelic equivalence (X-axis = 0, Y-axis = −1) and enhances exclusive maternal or paternal methylation (positive Y-axis values for X-axis = −1 or +1).

The Ref and Alt alleles were called in GM02639 MeDIP data as well and the Ref and Alt genotype ratios for CT were plotted against those obtained from KD. We found that there were large scale allelic imbalance represented in the genotype distributions (Fig 5E). The CT and KD genotype ratios were more anticorrelated in GM02639 (r^2^=0.000169) than HEK293T (r^2^=0.130321).

To objectively establish the extent of allelic imbalance in MeDIP between CT and KD, we sequenced the paternal and maternal DNA for GM02639 (see methods for details). We fished out only those regions at which the two parents were homozygous for different alleles such that at these loci the GM02639 could only be heterozygous in the absence of any sweeping allelic imbalances in methylation. Out of 11526 such loci, GM02639 was expected to be heterozygous with Mat^Alt^/Pat^Ref^ genotype for 6613 loci and Mat^Ref^/Pat^Alt^ genotype at 4913 loci. We set an arbitrary threshold of Diff/Sum ratio such that values ranging between −0.5 and 0.5 were regarded as heterozygous (green shaded region of the scatter; Fig 5F). In this case, the heterozygosity represented biallelic methylation within the range of Diff/Sum ratio threshold. Four types of unexpected deviations from the expected heterozygosity were observed in both CT and KD (non-green shaded regions of the scatter; Fig 5F). These were as follows: Mat/- or -/Pat (due to a monoallelic methylation bias similarly in CT and KD; red shade in Fig 5F), Mat/- or -/Pat in CT and Mat/Pat in KD (due to a loss of monoallelic methylation bias KD; aqua shade in Fig 5F), Mat/- or -/Pat in KD and Mat/Pat in CT (due to a gain of monoallelic methylation bias KD; purple shade in Fig 5F), and an allelic flip from Mat/- in CT to -/Pat in KD and from -/Pat in CT to Mat/- in KD (due to a random allelic choice for methylation; yellow shade in Fig 5F). The loci falling in the purple, aqua and yellow shaded regions of the figure 5F represent regions at which cytosine methylation occurs with a stochastic allelic bias that depends on CGGBP1. The red-shaded regions represent a stochastic deviation from heterozygosity independent of CGGBP1 resulting in correlated parent-of-origin identities between CT and KD (Fig 5F). To quantify the effect of CGGBP1 depletion on stochastic allelic choices and resulting deviations from heterozygosity, we calculated the differences in distribution of alleles with parental identities between CT and KD. As shown in figure 5G, the extreme allelic bias with a completely monoallelic methylation for the maternal as well as the paternal alleles was increased after CGGBP1 depletion with a concomitant loss of biallelic methylation similarly for the maternal or the paternal alleles. The CTCF-binding sites were not associated with any specific type of allelic bias (not shown). These results confirmed that CGGBP1 depletion exacerbates the allelic imbalance in cytosine methylation in a stochastic manner giving rise to maternal or paternal biases in methylation as well as an allelic reversal of methylation. The cytosine methylation represented in the CT data itself has a considerable allelic imbalance and that the stochastic change in cytosine methylation caused by CGGBP1 depletion further enhances the extent of allelic imbalance.

## DISCUSSION

In this study, we have assayed the effects of CGGBP1 depletion on global patterns of cytosine methylation in two different cell types using MeDIP. Unlike WGBS, MeDIP provides a lower resolution information but offers an advantage in alignment frequency of sequence reads generating a higher effective coverage per locus than WGBS. The MeDIP also allows querying of the MeDIP-seq data for sequence patterns and motifs with a higher confidence than WGBS. Using WGBS in 1064Sk cells, we have previously reported that the methylation changes caused by CGGBP1 depletion are a near random combination of GoM and LoM [27,28]. The deductions of gain or loss of methylation in a typical WGBS assay are confounded by two related reasons: (i) the WGBS technique captures and reports unmethylated as well as methylated cytosines without any weightage of the density of methylated or unmethylated cytosines [32], and (ii) the Bayesian probability frameworks in which cytosine methylation changes are called from a WGBS experiment rely on local cytosine and methylcytosine densities [33–36]. Thus, the large fraction of cytosines, that remains unmethylated or retains methylation between CT and KD fibroblasts in WGBS assays, confounds the distinction between deterministic or stochastic changes in methylation. In this study, to better understand the mechanisms of cytosine methylation regulation by CGGBP1, we employed MeDIP as a complement to our previous WGBS approaches [27,28]. Unlike WGBS, MeDIP-seq sacrifices the base level resolution of methylation information for a semiquantitative estimation of methylation. The length resolution of methylation signal in MeDIP-seq is governed by the size of the input DNA fragments which is also reflected in the mean read lengths. The representation of a region in the MeDIP-seq thus depends on local methylcytosine density in a particular region and the frequency with which it is captured in MeDIP. These parameters are expected to vary intracellularly between alleles as well as due to intercellular heterogeneity in cytosine methylation. A thorough analysis of the MeDIP-seq data from CT and KD allows us to measure stochastic or deterministic quantitative changes in cytosine methylation over short sequences. In this case, we restricted our analyses to a minimum resolution of 0.2 Kb (a convenient bin size that is larger than the mean length of the sequence reads). For a comparison, we have analyzed MeDIP-seq in CT and KD samples of foreskin fibroblasts (a cell type in which CGGBP1-regulated methylation has been studies earlier) and HEK293T (cells which are less sensitive to CGGBP1 depletion than fibroblasts). Moreover, in HEK293T cells we have recently shown that CGGBP1 determines the chromatin occupancy of CTCF at repeats versus motifs [20] and the current findings of methylation change in KD allow us to interpret cytosine methylation as a possible means through which CGGBP1 regulates CTCF occupancy. We have analyzed the nature of methylation changes in the two cell types and characterized the stochasticity of methylation changes caused by CGGBP1 depletion. We have also combined a blind TFBS analysis with our prior knowledge of the CTCF-CGGBP1 axis to extract subtle but deterministic methylation changes at CGGBP1-regulated the CTCF binding sites that are RFM.

The equal and mirroring patterns of GoM and LoM in HEK293T CT and KD are explainable as an outcome of stochastic changes in methylation. A wide range of methylation differences between HEK293T CT and KD corresponded to a similar high level of entropy. A non-functional TP53 in stem cells induces *de novo* methyltransferases resulting in high global methylation that is less prone to decrease after prolonged culturing [37,38]. The SV40 T antigen-mediated inactivation of functional TP53 in HEK293T is thus expected to have a much higher buffering capacity against methylation changes. The relatively less dependence of HEK293T on CGGBP1 and the higher levels of methylation observed in it could thus be explained by stochastic nature of methylation changes caused by CGGBP1 depletion. The absence of TP53 in HEK293T could further augment the stochasticity of methylation and dilute any deterministic changes. In the light of our recent findings that CGGBP1 is required for proper CTCF occupancy on the chromatin in HEK293T [20], the CGGBP1-regulated CTCF binding sites are strong candidate regions where a quantitative change could be expected between CT and KD. We reason that the high level of methylation stochasticity in HEK293T preclude the detection of methylation changes at CTCF binding sites. As a corollary, in fibroblasts, which are very sensitive to CGGBP1 depletion, do not have aberrant TP53-driven *de novo* methylation activity and may loose methylation further upon prolonged culturing, less entropy was observed in methylation changes. Supporting this, we observed that the stochasticity in fibroblasts was higher in regions with weaker methylation change. The regions with stronger change in methylation between CT and KD showed more deterministic change. The PCA accordingly revealed that RFM were the drivers of this deterministic change in methylation in fibroblasts. A higher GoM than LoM in GM02639 was different from the pattern seen in HEK293T. The directional change in cytosine methylation in GM02639 was detectable due to a lesser stochasticity than HEK293T, especially at higher levels of GoM. This deterministic change in methylation was concentrated at RFM. Interestingly, despite 90% knockdown of CGGBP1, HEK293T maintained strongly stochastic cytosine methylation in CT and KD, whereas just 50% knockdown in GM02639 caused a much less stochastic change in GM02639. This shows that there is a cell type specificity in the cytosine methylation changes caused by CGGBP1 knockdown and it is consistent with the previous findings [20] that HEK293T express higher levels of CGGBP1 and show less dependence on it as compared to fibroblasts. The stochasticity is observed in a large population sum total of methylation signals derived from pools of cells with mosaic methylation patterns.

Primary cells in culture rapidly lose methylation whereas immortalized cells are resistant to rapid changes in methylation under the same conditions. In our experiments we also see that the net quantitative change of methylation is close to zero in HEK293T cells whereas the fibroblasts show a much stronger net increase. The very high entropy of methylation patterns in HEK293T CT and KD compared to those in GM02639 are difficult to explain through random intercellular variations of methylation patterns as unlike WGBS data, we can not deconvolute [39] the MeDIP data to predict the cellular heterogeneity in the two cell cultures and any differences between them. Given that CGGBP1 depletion slows down or arrests cell cycle [40–42], the intercellular heterogeneity is expected to remain unaffected or diminish in KD as compared to CT. These facts suggest that a major fraction of stochasticity in methylation patterns, that is retained after CGGBP1 depletion, is due to factors other than intercellular heterogeneity. There is overwhelming evidence that the stochasticity of methylation patterns are due to localized allelic imbalances in methylation [3,43–46]. We tested the possibility that between CT and KD the stochastic changes in methylation patterns are due to random inter-allelic differences in methylation levels. Indeed we found that in both the cell types there were unexpectedly high levels of allelic imbalances in methylation. It was intriguing that between CT and KD, the allelic imbalances were qualitatively different. This included mostly a gain or loss of monoallelic or biallelic methylation. A significant subset however showed a highly unexpected monoallelic swap between the two parental genotypes in methylation between CT and KD. The allelic switch of methylation requires a stochasticity in methylation that is very dynamic and highly entropic. Since methylation as well as CTCF binding are known to be key regulators of genomic imprinting [8,13,47], we needed to rule out the possibility of a non-random allelic choice of methylation upon CGGBP1 depletion. To ensure that there was no parent of origin bias in the allelic switch between CT and KD, we sequenced and characterized the parental DNA of the fibroblasts. The allelic imbalance analysis with parent of origins defined convincingly demonstrated that the allelic imbalance of methylation in CT and KD are not biased towards any parent of origin and thus highly stochastic in nature.

Previously, WGBS has shown that the CpG methylation increases as well as decreases at repeats although the majority of methylation is at CHG and CHH cytosines [27]. Including all the three contexts, the prevalence of CHH cytosine methylation leads to the identification of G/C skew as a signature of sequences showing an increase as well as decrease due to CGGBP1 depletion [28]. The current MeDIP seq characterizes methylation patterns in a context independent manner and only much stronger differences in methylation (than those characterized through WGBS) would be able to affect the enrichment with methylcytosine antibody. Thus, even under high levels of stochasticity, the identification of quantitative differences in MeDIP signals is a strategy that has allowed us identification of a panel of TFBSs as targets. The strength of our TFBS strategy is a search for motifs that occur commonly in two disparate cell types used in this study. This ensured that the motif discovery was not due to cell type specific stochasticity in the methylation patterns but just due to the differences between CT and KD. This approach is prone to false negatives but robust against false positive motif detections. Binding of some TFs to their target sequences determines methylation turnover by regulating the access of methylation machinery to the DNA [48]. This is known for CGGBP1 [25] and its depletion can thus lead to a random in methylation levels. In addition, methylation regulation by CTCF, REST, SP1, EGR1 has been described [48–51] and in a double blind search we have found these factors to be significantly enriched in the differentially methylated genomic bins between CT and KD. Thus the TFBS commonly identified in the differentially methylated bins of HEK293T as well as fibroblasts lead us to identify functionally relevant non-stochastic methylation changes caused by CGGBP1 depletion. The DNA-binding of TFs with binding site overrepresentation in differentially methylated bins are regulated by cytosine methylation as well as regulate cytosine methylation. Of all the TFs identified as significantly overrepresented in the CT KD DMRs, CTCF is the one for which a regulation by CGGBP1 has been demonstrated [20]. One of the proposed mechanisms of methylation regulation by CTCF, consistent with the stochasticity of methylation changes observed between CT and KD, is that CTCF sterically hinders methylation machinery access to the DNA [52]. We thus followed up the CGGBP1-regulated CTCF-binding sites to extract regions of non-stochastic changes. We have reported that CGGBP1 regulation of CTCF occupancy at repeats and motifs are inversely related [20]. There is no evidence that CTCF-repeat interaction is indirect and thus different from the direct binding of CTCF to its motif. Thus, the methylation sensitivity of CTCF binding at motifs may not necessarily apply to CTCF binding at repeats. This is supported by the findings that L1 repeats are not differentially represented in differentially methylated regions, but the CTCF motifs are specifically differentially methylated between CT and KD. Following up on this lead, we segregated the CTCF binding sites as RFM and MFR and confirmed that CGGBP1 regulated methylation changes affect CTCF-binding motifs and not the repeats non-stochastically. Our findings suggest that different cell types show different types of methylation changes at CTCF motifs upon CGGBP1 depletion. These results also suggest that motif-specific methylation change may be a mechanism underlying the shift in CTCF binding preferences from repeats to motifs upon CGGBP1 depletion.

Stochasticity is an innate property of cytosine methylation and has been addressed in multiple studies [43,53,54]. In our investigation here, the stochastic nature of methylation has remained dominant over the effects of CGGBP1 depletion on global methylation patterns. The fundamental question of why CGGBP1 depletion leads to such a widespread resetting of methylation remains unknown. We postulate that the widespread occupancy of CGGBP1 on the genome in the presence of normal amounts of CGGBP1 maintains a state of dynamic equillibrium wherein the CGGBP1-bound DNA remains unavailable for binding and activity of the methylation regulatory apparatus. The lowering of CGGBP1 levels disrupts this equilibrium such that the DNA denuded of CGGBP1 becomes more amenable to activity by the methylation regulatory apparatus.

Our results suggest an interplay between a stochastic disruption in methylation caused by CGGBP1 depletion and its effects on specific TFBSs, including CTCF. The disruption of methylation upon CGGBP1 depletion targets sites at which CTCF binding has been recently demonstrated. The cause-consequence relationship between methylation changes and CTCF binding at RFM remains unknown, but it is highly likely that it is a two-way feedback process. However, the disruption in CTCF occupancy at RFM seems to be functionally relevant as the methylation asymmetry in the flanks of these CTCF-binding sites seem to be affected specifically. These results complement our previous findings that the H3K9me3 signals exhibit asymmetry in the flanks on CGGBP1-regulated CTCF-binding repeats. It appears that CGGBP1 stabilizes a cytosine methylation profile at RFM that allows CTCF to maintain a barrier against methylation spread across the RFM. Thus, cytosine methylation homeostasis is a crucial entity at the interface of regulation of CTCF barrier activities by CGGBP1.

## CONCLUSION

Cytosine methylation is stochastic in HEK293T and GM02639 cells. CGGBP1 depletion in these cells changes cytosine methylation patterns such that the stochasticity remains unperturbed but the allelic imbalances that underlie the stochasticity are different in CT and KD. Embedded in the largely stochastic methylation patterns in CT and KD are specific TFBSs which show a cell type-specific quantitative change in methylation. One of these TFs is CTCF. The methylation patterns at CTCF-binding MFR and RFM show different dependence on CGGBP1. The MFR methylation remains stochastic between CT and KD whereas the RFM shows a non-stochastic change. The methylation changes between CT and KD were stochastic with no parent-of-origin bias. The non-stochastic methylation changes at RFM were not due to lower levels of stochastic allelic imbalances.

## LIST OF ABBREVIATIONS

CT, KD, RFM, MFR, LoM, GoM.

## DECLARATIONS

### Ethics approval and consent to participate

Not applicable

### Consent for publication

Not applicable

### Availability of data and materials

An additional file contains details of the methods and materials used.

## AUTHORS’ CONTRIBUTIONS

MP, DP and SD performed the experiments. All authors participated in data analysis and manuscript writing. US supervised all aspects of the work. The manuscript is read and approved by all the authors.

## ACKNOWLEDGEMENTS

The authors acknowledge the Gujarat Biotechnology Research Centre for sequencing services, Coriell cell repository for fibroblasts (GM02640, GM02641 and GM02639), NCCS Pune for HEK293T cells and Mr. Sudeep N Banerjee (ISTF, IITGN) for help with computational resources.

## Methods

### Cell culture, transfections and lentiviral transductions

HEK293T (NCCS, Pune), and human foreskin fibroblasts from Coriell Cell Repository (Son = GM02639, Parents = GM02641 and GM02640) were cultured in DMEM (HiMedia or HyClone) supplemented with 10% fetal bovine serum. HEK293T cells were subjected to lentiviral transduction with non-targeting shRNA (CT) and CGGBP1-targeting shRNA (KD) as described before [20]. The transduced cells were selected using Puromycin (10 µg/ml) for a week. The cells were lysed and genomic DNA was isolated using standard phenol:chloroform:isoamyl alcohol extraction method followed by ethanol precipitation and dissolution in 1xTris-EDTA buffer. For GM02639, the cells were transfected with non targeting siRNA (CT) or CGGBP1-targeting siRNA (KD) twice, once after 24h and second after 72h. The siRNA transfections were intended for a mild CGGBP1 dpeletion. The cells were collected and processed for genomic DNA isolation as described above for HEK293T cells. Genomic DNA was isolated from the parental fibroblasts GM02641 and GM02640 without any transfections or transductions. The siRNA CGGBP1-targeting (4392422, ThermoFisher scientific) for KD and non-targeting (4390844, ThermoFisher scientific) for CT were transfected with the Oligofectamine™ Transfection Reagent (12252011, ThermoFisher Scientific). The transfections were performed as per the manufacturer’s instructions.

### Western blotting

The knockdown of CGGBP1 was confirmed by performing Western blots on the lysates of transduced HEK293T and transfected GM02639 cells at 96h. The primary antibodies were a Rabbit anti-human CGGBP1 polyclonal (10716-1-AP, Proteintech) or Mouse anti-human GAPDH monoclonal (MA5-15738, Invitrogen). The secondary antibody was HRP conjugated Goat anti-Rabbit (NA934, GE Life Sciences). The samples were resolved on 4-12% Bis-Tris NuPAGE (Invitrogen) gels, transferred to PVDF membranes (3010040001, Merck), blocked for 1h in blocking buffer (5% dry milk v/w and fetal calf serum 10%v/v (HiMedia) in 1x TBST buffer) followed by overnight incubation with the primary antibody overnight at 4°C (1:100 dilution in 1x TBST buffer). Membranes were washed in 1x TBST, incubated with HRP conjugated anti-rabbit secondary antibody (1:5000 dilution in blocking buffer) for 2 hours at room temperature followed by washing with 1xTBST. The signals were developed using ECL substrate (Pierce) and captured using chemiluminescence imaging setup (GE Life Sciences).

### Methyl(cytosine) DNA immunoprecipitation (MeDIP)

The DNA isolated from HEK293T and GM02639 cells were sonicated to obtain DNA fragments of size range 150-300 bp. The conditions for sonication were 30 cycles of 30 seconds ON and 30 seconds OFF (Bioruptor, Diagenode) with intermediate mixing. 20 µg DNA for each sample was used as input for MeDIP. DNA was incubated with 1x MeDIP master mix (10mM Sodium Phosphate Buffer, 0.14M NaCl and 0.05% TritonX-100) containing a 5-methylcytosine antibody cocktail (5µg; MABE146, SAB2702243; Sigma and NBP2-42813, Novus Biologicals) overnight with tumbling at 4°C. Pre-washed Protein G sepharose beads (17-0618-01, GE Healthcare) were added to the mix and incubated for 2 hours with tumbling mixing at 4°C. The beads were allowed to settle, collected by gentle centrifugation and gently washed with 1x IP buffer three times. Further the washed beads were incubated with Proteinase K (10ug/ml) in a protein digestion solution (50mM Tris-HCl (pH 8.0), 10mM EDTA (pH 8.0) and 0.5 % SDS) containing for about 2 hours at 56°C with occasional mixing. The immunoprecipitated DNA was collected by subjecting the mix to centrifugation and collecting the supernatant into new tubes. The MeDIP DNA was purified using the PCR cleanup kit (A1460, Promega).

### Sequencing of MeDIP and genomic DNA

The sequencing libraries for the MeDIP DNA (CT and KD from HEK293T and GM02639 cells) and genomic DNA (GM02641 and GM02640 cells) were generated according to protocol mentioned elsewhere [20]. The sequencing was done using the Ion Proton S5 sequencer.

### Mapping and quality control

The reads obtained post-sequencing was controlled for the low quality reads through filtering out the reads having lower than q20 threshold. The initial QC was controlled through fastq validation using fastq validator for filtration of any trimmed reads. The mean read length for all the samples was around 150 bp. The reads were mapped to repeat unmasked human genome hg38 with default mapping conditions using Bowtie2. Samtools was used for SAM to BAM conversions and sorting and indexing of the BAM file. Bedtools bedtobam and bamtobed functions were used for interconversions of BAM and BED files. The sequences from the reference genome (hg38 masked or unmasked) were obtained using bedtools getfasta option in bedtools. The fasta manipulations (format conversions, shuffle, statistics) were done using various functions in seqkit. The base composition and the cytosine contexts identification was done using the compseq function in the EMBOSS toolkit.

### Variant calling

The mapped reads obtained as BAM output were subjected to variant calling using bcftools. The BAM file was subjected to mpileup followed by the bcftools call to identify variants across the sequenced locations for each datasets. The variants were filtered for their minimum coverage of 5 reads for each sample.

### MeDIP signal at 0.2 Kb bins

Genome-wide 0.2 Kb bins were created through bedtools make-windows option. The MeDIP signal for all the samples at these bins was obtained by bedtools coverage option with minimum 50 percent of read overlapping with the bin. For pairwise analysis the bins were retained with a minimum signal of three reads for CT and KD combined. The pairwise signal comparisons by Diff/Sum ((KD-CT)/(KD+CT)) was done for the signals obtained from HEK293T and GM02639 cells respectively. The frequency distribution of these Diff/Sum values across these filtered bins was plotted to compare the methylation changes for HEK293T and GM02639 cells respectively.

### JASPAR-wide motif identification

The overlapping coordinates between bed files were generated using bedtools intersect. The 0.2 Kb bins were filtered for the threshold of |diff/sum| of more than 0.2 for HEK293T and GM02639 cells. These filtered bins were subjected to the motif identifications through fimo tool (MEME suite) for JASPAR wide motifs (519 in total) using the stringent threshold of 1E-6. In parallel, the unfiltered bins genome-wide were also subjected to the same analysis to generate expected background motif occurrence frequencies. The randomisation of the bed coordinates was done using bedtools shuffle option. These shuffled genomic coordinates were also subjected to motif finding using fimo. The comparison of the expected and observed motif frequencies was performed for each transcription factor individually using Chi-square test function in Graphpad Prism 8.

### Heatmap on MeDIP signal

The MeDIP signal was plotted on the bins for the filtered motifs as a Diff/Sum of average signal for CT and KD for each transcription factor. The plotting was done using R package ComplexHeatmap tool using the single clustering method.

### Heatmaps and average summary profile

The bigwig for the methylation signal was generated by using bamCoverage tools from deeptools. The scaling factor was applied to normalize the total readout of CT and KD samples for GM02639 cells. No scaling factor was applied for generation of bigwigs in CT and KD for HEK293T cells. The methylation signals were plotted as heatmaps using the deeptools plotHeatmap. The average summary plots along with standard deviation for methylation signals were plotted using plotProfile function with plotType std option in deeptools. The matrix used for these functions were generated using deeptools computeMatrix function.

### Correlation analysis

Correlation between MeDIP signals was performed by using multiBigwigSummary tool from deeptools. Methylation signals were compared at bin sizes of 10 Kb, 5 Kb, 1 Kb and 0.2 Kb. Correlation between samples was calculated by Spearman method by using deeptools plotCorrelation tool.

### PCA analysis

The matrices obtained from the multiBigwigSummary were subjected to PCA analysis using plotPCA function in deeptools.

### Repeat content analyses

The repeat-masked and unmasked human genome (hg38) were used from the UCSC genome browser. Locally installed version (version open-4.0.9) of Repeat-masker was used for repeat-masking. RMBlast (NCBI) was used as a repeat search engine with and the repeat database used was Repbase (version available in 2018).

### CTCF binding site identification and motif finding

The peaks were celled on the CTCF reads [20] that entirely overlapped with the methylated regions, across the methylation bins using MACS2.0 callpeak. *De novo* motif search was carried out by locally installed version of MEME suite (version 5.0.3) tools meme. The motif search in GoM, LoM and no change 0.2 Kb bins were performed by using default option with motif length 19 (-w value of 19). MEME suite tool fimo was used to find predicted motifs in sequences. Predicted motifs were found with a default threshold of 1E-4.

### Statistical analyses, graphing and genome browser viewing

Statistical analysis was performed by using Prism version 8 (GraphPad) on numerical data stored and sorted in OpenOffice Spreadsheet. Visualisation of MeDIP signal at genomic regions were carried out by locally installed Integrated Genome Viewer.

### Restriction digestion of genomic DNA for qPCR-based methylation detection

Genomic DNA was isolated from human dermal fibroblasts (106-05A,Sigma) transfected with Dharmacon siRNA cocktails as follows: Non-targeting (D-001910-10-20, Dharmacon) designated as CT and KD CGGBP1-targeting (E-015703-00-0020, Dharmacon) designated as KD. Genomic DNA was extracted as described above for MeDIP. DNA was sonicated to mean length of 1-1.5 Kb and subjected to restriction digestion. CT and KD DNA were subjected to restriction digestion by methylation-dependent or methylation sensitive endonucleases. The digested DNA were used as templates for quantitative PCR for a panel of candidate regions (Table S4). The Ct values obtained from Cytosine methylation (all contexts)-dependent digestion using *Mcr*BC were normalized against undigested input. Methylation sensitive digestions were performed using *Hpa*II and the Ct values were normalized using corresponding *Msp*I digestions. Following enzymes were used: *Hpa*II (R0171S, NEB), *Msp*I (R0106S, NEB) and *Mcr*BC (M0272S, NEB). For each restriction enzyme digestion, 1 µg of DNA template was used with 2 µl of enzyme for 6h at 37°C. The digested DNA was used as a template for qPCR using SYBR-Green PCR 2x mix (1725124, Bio-Rad), InstaQ96 (HiMedia) and the various primers as indicated in Table S4. Relative quantitative changes were calculated using delta-delta Ct method.

