## Supplementary material for "CGGBP1-regulated cytosine methylation at CTCF-binding motifs resists stochasticity": Legends to supplementary figures and tables

Figure S1: CGGBP1 depletion in HEK293Tcells.
HEK293T cells were transduced with non-targeting shRNA or CGGBP1-targeting shRNA lentiviruses. Lentivirus-transduced cells were selected with 10 µg/ml Puromycin for 1 week and subjected to immunoblotting. The level of CGGBP1 and GAPDH are shown in upper and lower panel respectively. A CGGBP1 knockdown of approximately 77% is observed when normalized to GAPDH levels.

Figure S2: Methylation differences between CT and KD are discernible at small genomic length ranges. Genome-wide methylation signal distribution was compared between CT and KD by using "deeptools multiBigwigSummary". Methylation signals were compared at bin sizes of 10 Kb, 5 Kb, 1 Kb and 0.2 Kb. Correlation between CT and KD was computed by Spearman method by using "deeptools plotCorrelation" for HEK293T cells.

Figure S3: CGGBP1 depletion in GM02639 cells.
GM02639 cells were transfected with non-targeting or CGGBP1-targeting siRNA twice at 24 and 72 hours post-seeding. Cells were harvested at 96 hours. Immunoblotting results for CGGBP1 (upper panel) and GAPDH (lower panel) show approximately 55% knockdown of CGGBP1 when normalized to the level of GAPDH.

Figure S4: Methylation differences between CT and KD are discernible at small genomic length ranges. Genome-wide methylation signal distribution was compared between CT and KD by using "deeptools multiBigwigSummary". Methylation signals were compared at bin sizes of 10 Kb, 5 Kb, 1 Kb and 0.2 Kb. Correlation between CT and KD was computed by Spearman method by using "deeptools plotCorrelation" for GM02639 cells. These correlation coefficients can be compared with those for HEK293T (Fig S3).

Figure S5: Repeat content analysis in HEK293T CT and KD MeDIP DNA shows subfamily-specific methylation changes. Methylation bin frequency plots for HEK293T CT and KD. CT and KD reads for each methylation bin (From 5 to 30) were merged and sequences for merged regions (>150 bp long) were extracted and subjected to repeat identification. The figures depict the occurrence of the three most populous repeats (Satellites, L1-LINEs and Alu-SINEs). **A:** No overall differences in repeat content were observed between CT and KD. **B:** A classification of the Alu SINEs into J, S and Y subfamilies revealed subfamility-specific differences in methylation between CT and KD. AluJb and AluSx showed consistently higher methylation in KD across all the methylation bins (top two panels), while AluY showed lower methylation in KD (bottom panel). **C:** A subfamily classification of L1 repeats revealed that the L1HS and P family LINE1 such as L1P1, L1P2 and L1PA4 are reduced in KD although these repeat subtypes are more prevalent in highly methylated regions. **D:** In contrast, the early originated LINE1 such as L1M1, L1M3, L1M4 and L1M5 showed increased methylation in KD and these repeat subtypes are prevalent in regions with low levels of methylation.

Figure S6: Repeat content analysis in GM02639 CT and KD MeDIP DNA shows subtle subfamily-specific methylation changes. CT and KD MeDIP repeat identification was performed as described above for Figure S3. repeat identification and the occurrence of the three most populous repeats (Satellites, L1-LINEs and Alu-SINEs) were analyzed. **A:** Satellite and L1 repeats were overrepresented and underrepresented respectively in KD. No difference in Alu-SINEs content was observed however. **B:** Unlike HEK293T data, the subfamily classification of Alus does not reveal any subfamily-specific methylation differences between CT and KD in GM02639. **C** and **D:** L1 repeat subfamily classification showed no consistent differences in methylation between CT and KD.

Figure S7: Highly similar CTCF binding motifs are present in regions undergoing GoM, LoM or showing no methylation change upon CGGBP1 depletion. Methylation signal for CT and KD was calculated for each 0.2 Kb bin for HEK293T and bins were grouped into GoM, LoM and "No change" (as described in method in details). CTCF motif positive 0.2 Kb bins were filtered out from each group. Motif positive GoM, LoM and "No change" 0.2 Kb bin sequences were subjected to *de novo* motif search by using MEME suite.

Figure S8. Quantitative PCR (double delta analysis of relative changes in levels of methylated DNA) on CT and KD DNA from GM02639 shows widespread differences between CT and KD. **A:** qPCR on *Mcr*BC-digested DNA (digests DNA flanking methylcytosine) shows a significant ($p<0.05$, n=3 technical replicates, unpaired T test) loss of methylation at genomic regions representing the PEG10 locus, GRB10 locus and a CpG island termed CpG-16. Conversely, a loci representing TDG, CTCF-binding site termed CTCF-2-2, TET3 and NAP1L5 showed a gain of methylation. **B:** qPCRs on *Mcr*BC-digested CT and KD DNA for multiple other regions showed erratic methylation disturbances that were insignificant. **C:** A *Hpa*II digestion qPCR showed an increase in methylation at CpG sites for multiple loci including Alu repeats and CTCF-binding sites ($p<0.05$, n=3 technical replicates, unpaired T test). All *Hpa*II Ct values were normalized against corresponding Ct values obtained after *Msp*I digestions. **D:** Widespread CpG methylation disturbances (with no significance) were also observed at multiple other loci. Refer to table S4 for exact location and PCR details.

**Supplementary Tables**

Table S1. Sequencing and alignment statistics for CT and KD MeDIP in HEK293T and GM02639.

Table S2. GC content and cytosine context representation for CT and KD MeDIP in HEK293T and GM02639.

Table S3. The MeDIP signal correlation between CT and KD at different genomic bin sizes decline with a reduction in bin size specifically as randomization of coordinates (and thus corresponding sequences) changes the correlation stochastically away from the observed correlation coefficients with actual MeDIP sequences (refer to Figure S2 for a comparison with correlation coefficients without any randomization).

Table S4. Primer names, locations, sequences and annealing temperatures used for the candidate region methylation analyses shown in Figure S6. The cyclic denaturation (95°C, 30 seconds) and extension (70°C) steps were the same for all primer combinations. Data were collected post-extension at 80°C as mentioned in methods.
