## Supplementary figures and images for "CGGBP1-regulated cytosine methylation at CTCF-binding motifs resists stochasticity"

### Supplementary Figures

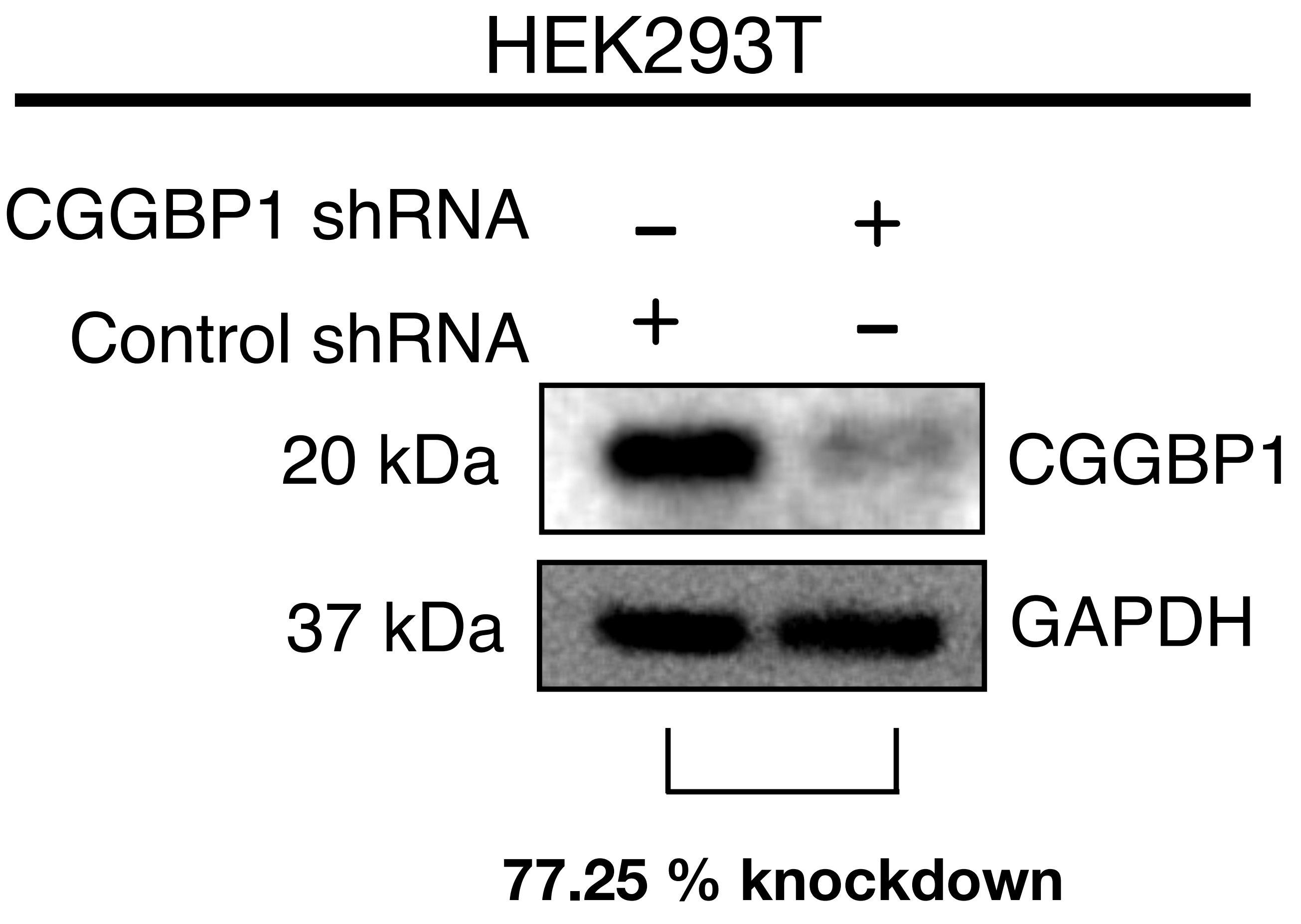

Figure S2

HEK293T

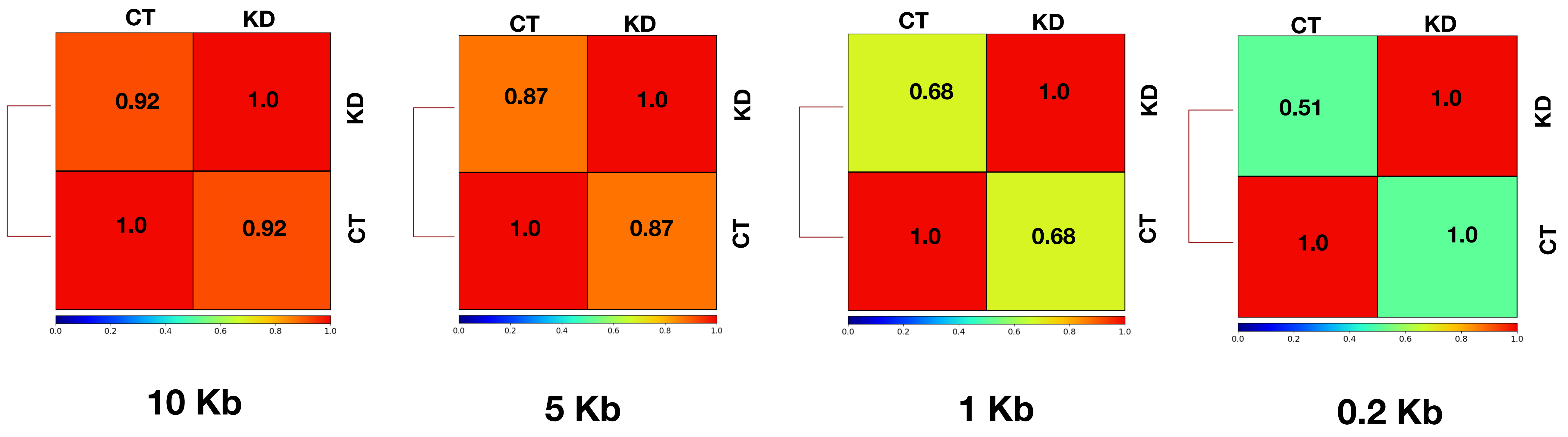

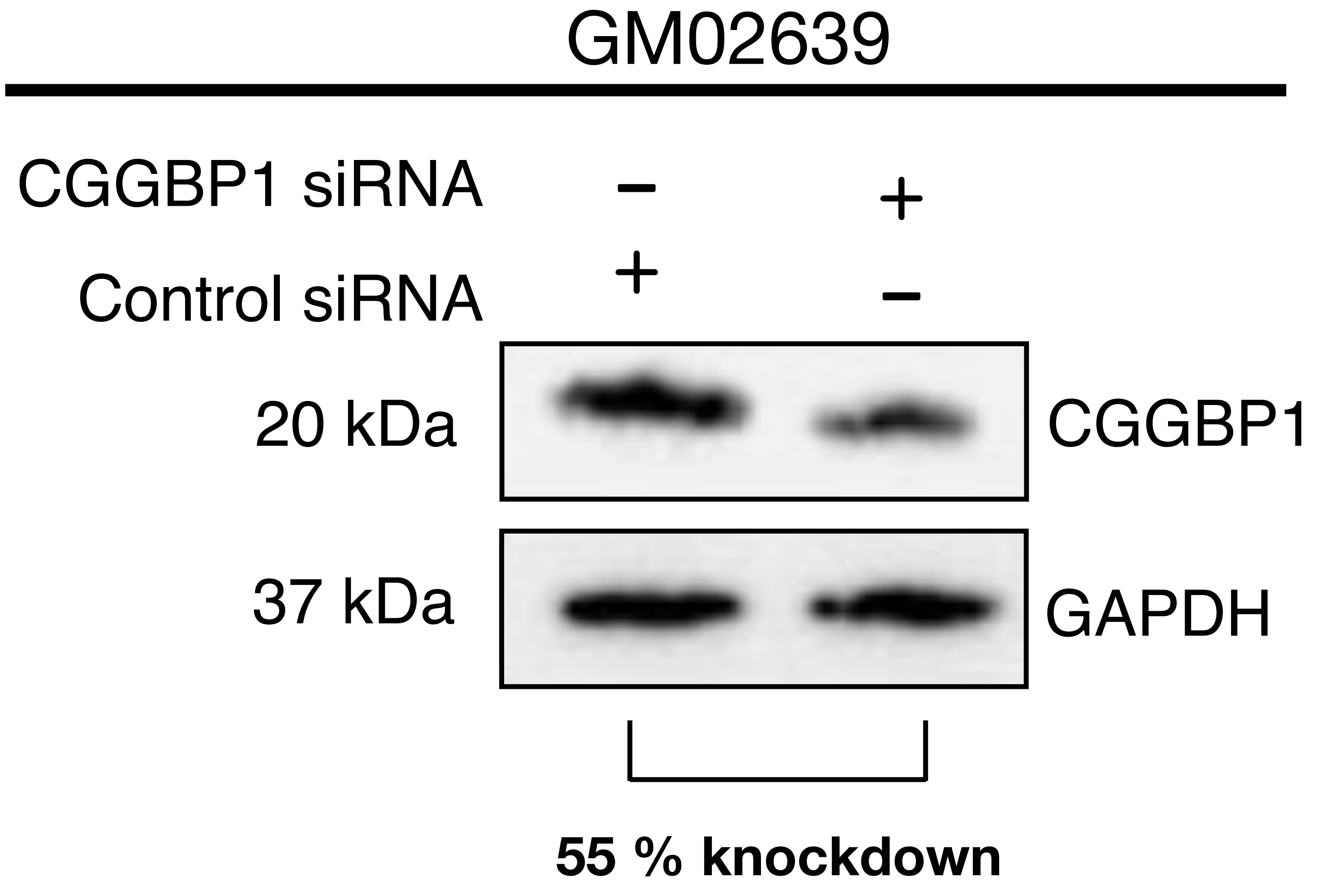

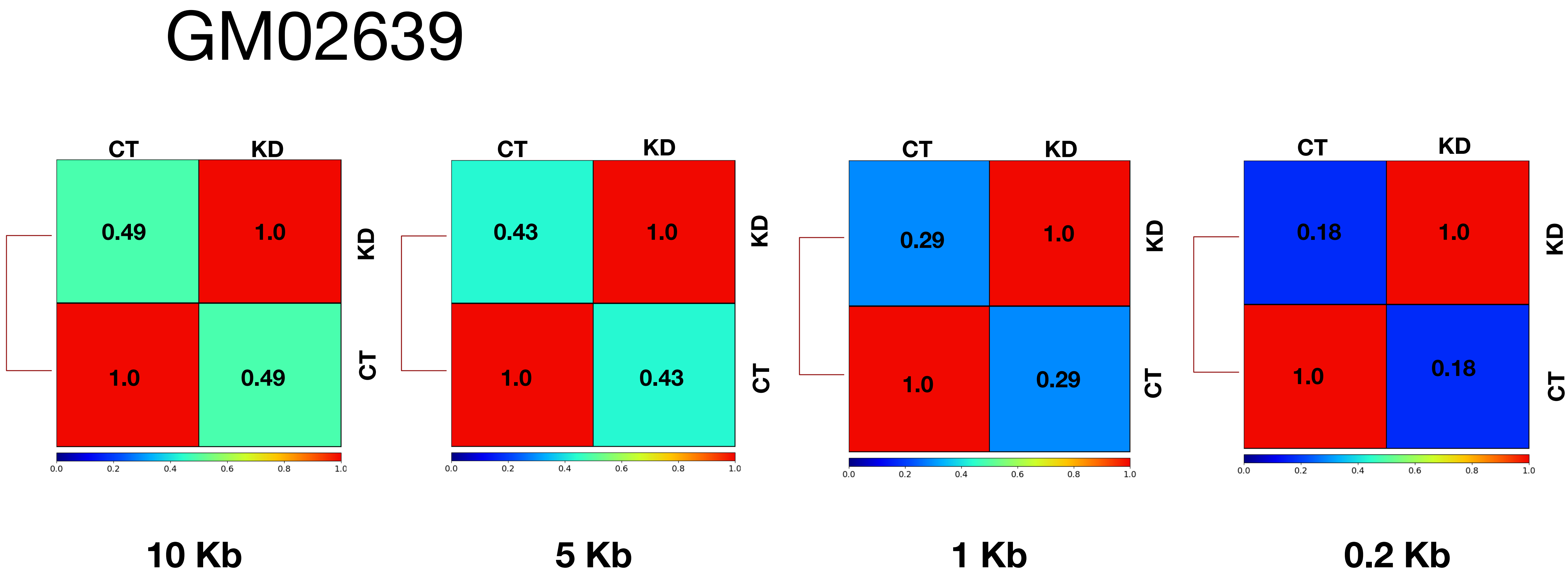

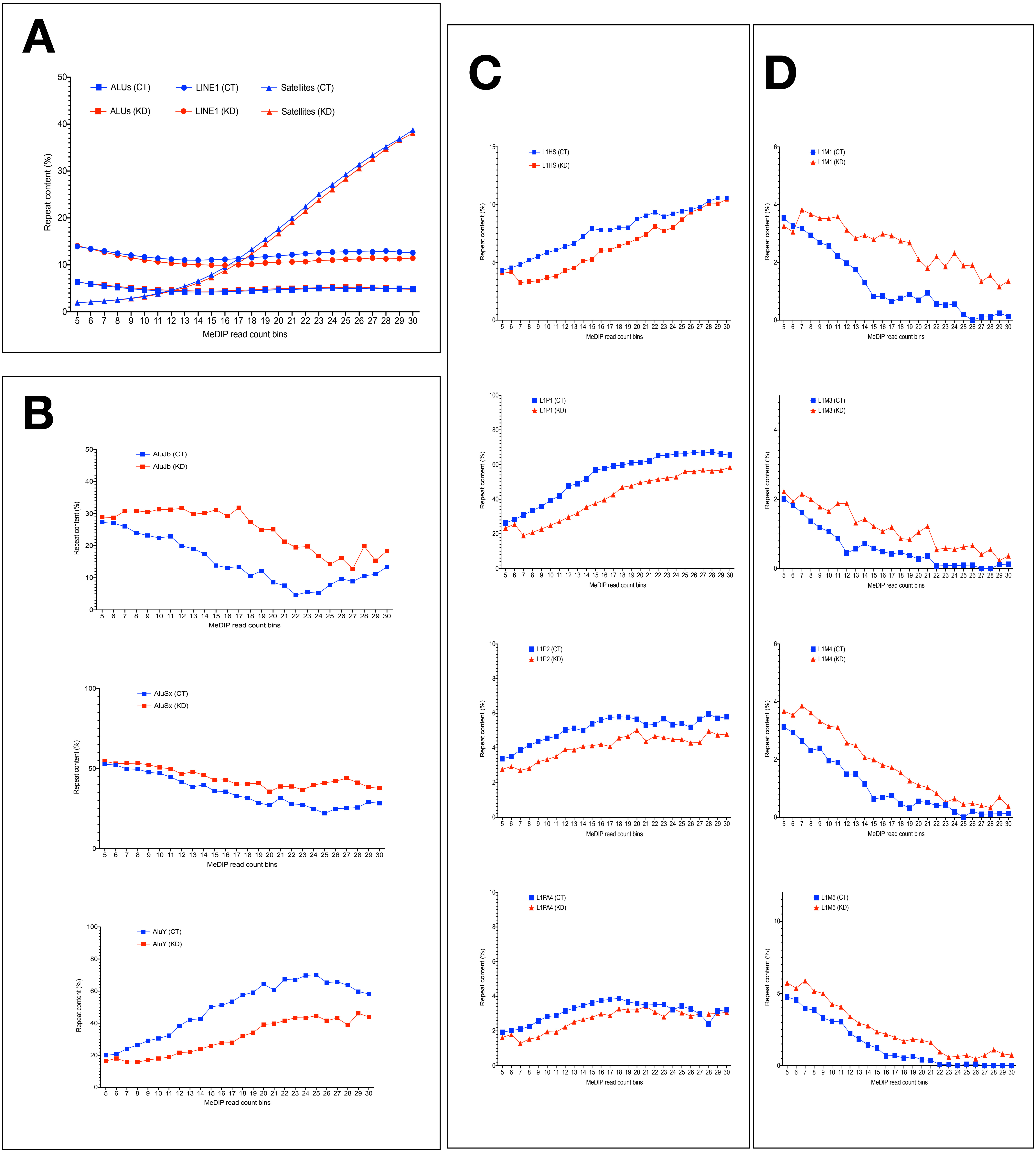

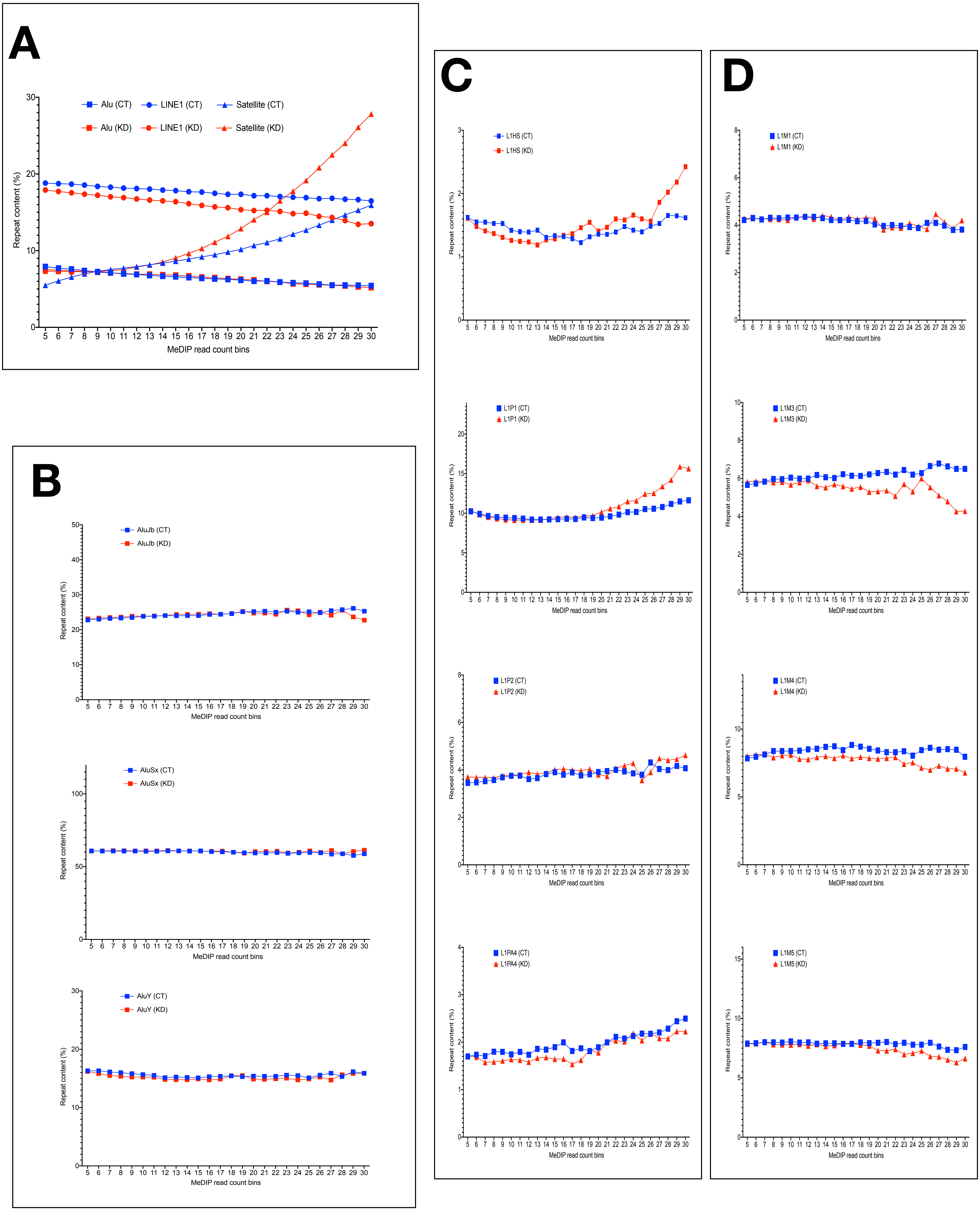

Figure S7

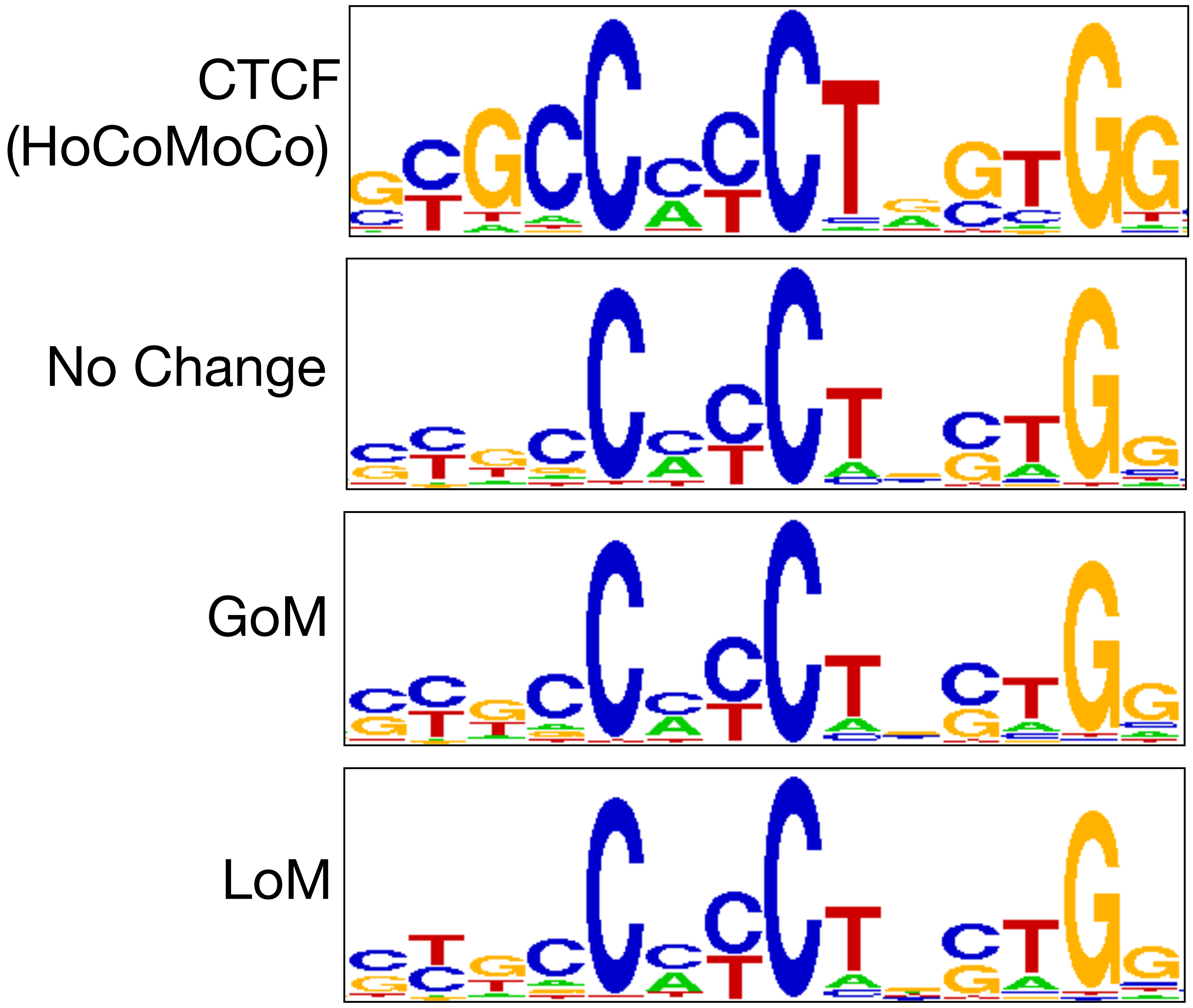

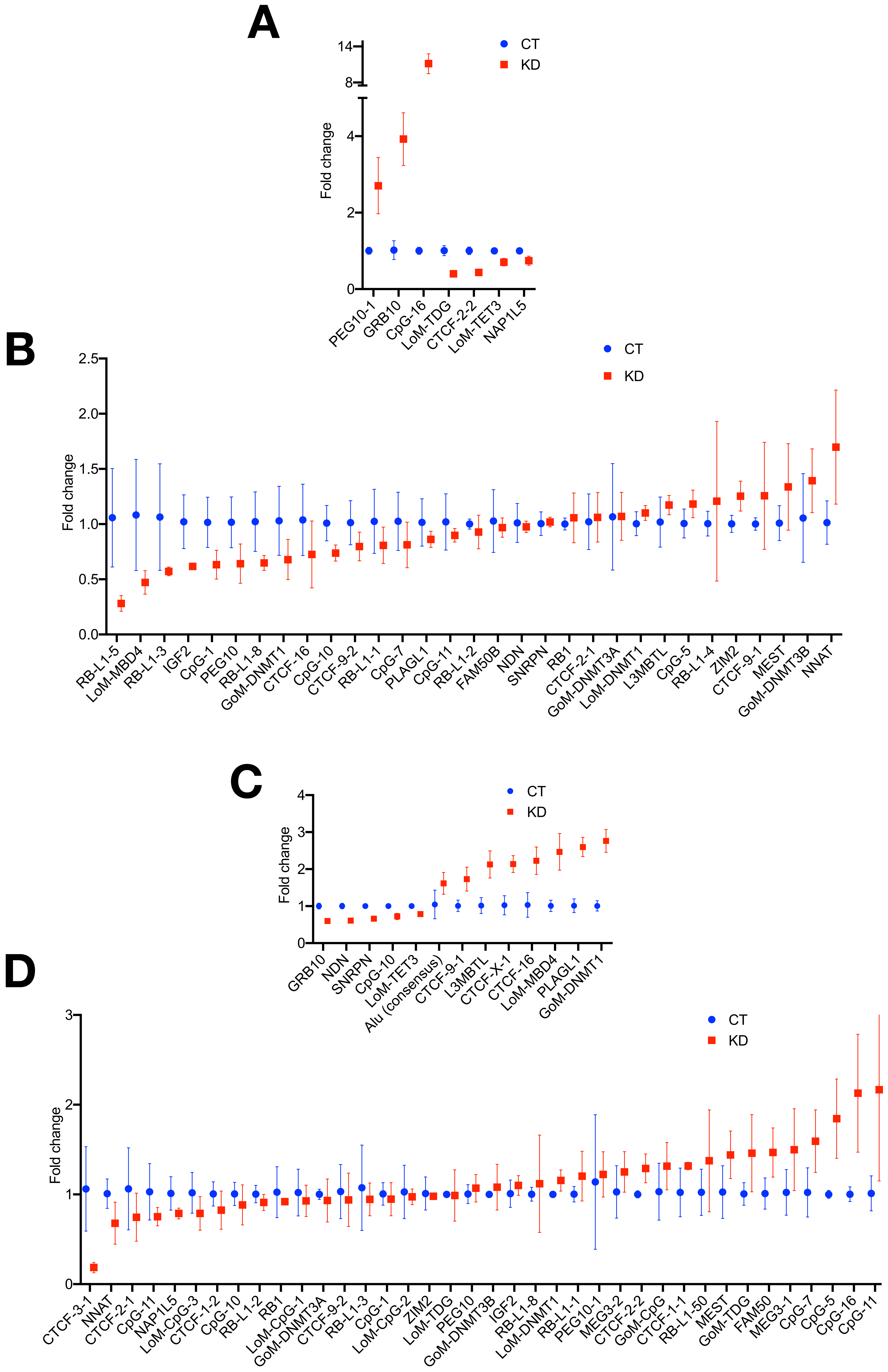
