## Supplementary Tables for "CGGBP1-regulated cytosine methylation at CTCF-binding motifs resists stochasticity"

Table S1

| Name | Total reads | Total mapped reads | Total unmapped reads | Reads % mapped | Reads % unmapped |
|---|---|---|---|---|---|
| HEK293T CT | 75441659 | 70195180 | 5246479 | 93.046 | 6.954 |
| HEK293T KD | 76069183 | 69404104 | 6665079 | 91.238 | 8.762 |
| GM02639 CT | 118197258 | 94073184 | 24124074 | 79.59 | 20.41 |
| GM02639 KD | 83791601 | 45964460 | 37827141 | 54.856 | 45.144 |
| GM02641 | 71104585 | 68684768 | 2419817 | 96.597 | 3.403 |
| GM02640 | 96230084 | 92852408 | 3377676 | 96.49 | 3.51 |

Table S2

| Samples | GC % | CpG % | CHG % | CHH % |
|---|---|---|---|---|
| HEK293T CT | 42.21 | 5.42 | 21.51 | 73.07 |
| HEK293T KD | 42.49 | 5.67 | 21.64 | 72.69 |
| GM02639 CT | 40.47 | 7.80 | 21.00 | 71.20 |
| GM02639 KD | 40.87 | 13.45 | 21.12 | 65.44 |

Table S3

| | | | | | | | |
|---|---|---|---|---|---|---|---|
| **10 Kb bins (Spearman r values)** | | | | | | | |
| CT::KD | 0.92 | CT shuffled::KD | 0.38 | KD shuffled::CT | 0.39 | CT shuffled:: KD shuffled | 0.78 |
| **5 Kb bins (Spearman r values)** | | | | | | | |
| CT::KD | 0.87 | CT shuffled::KD | 0.35 | KD shuffled::CT | 0.36 | CT shuffled:: KD shuffled | 0.69 |
| **0.2 Kb bins (Spearman r values)** | | | | | | | |
| CT::KD | 0.51 | CT shuffled::KD | 0.15 | KD shuffled::CT | 0.15 | CT shuffled:: KD shuffled | 0.21 |

Table S4

| TEMPLATE | LEFT PRIMER | RIGHT PRIMER | ANNEALING TEMPERATURE |
|---|---|---|---|
| ALU | GAGGCTGAGGCAGGAGAATCG | CGCCCAGGCTGGAGTGCAGTGGCGCG | 55 |
| CPG-1 | CGAGAGCACTACGCAGTCAG | TCTGACACCTAAGCCCTACCA | 53.5 |
| CPG-11 | ATTGCCTCACCTGGGAAG | GAGATTCCGTGGGCGTAG | 54.5 |
| CPG-10 | CGGGCCAGTGACAAAGAG | GCCATGGAGTCCTACGATGT | 54.5 |
| CPG-16 | TGGGACCTAGAGAACCGAGA | ATTGAGACATCAGCGGCATT | 54 |
| CPG-5 | GTGCCCAGGTAGAAGCAGAG | CGTCTCCATGTGCTGCTTT | 54 |
| CPG-7 | AGAACAGCGATTCCTTCGAG | AGTCCCTCGGCCAGTTTATC | 54 |
| CTCF-1-1 | TGCATCTGCAGAGAAGGAGA | AGCGAGACATACGCAGACCT | 55 |
| CTCF-1-2 | TGGTACTGCACCACTCTGGA | GAAAGTCCTAGCGGATCTGG | 55 |
| CTCF-16 | AGGTGCAGGGAATAATGCAG | TGCTTCCAGACATGGGTATG | 54 |
| CTCF-2-2 | CACACGTTTGGTGGCTTAAA | AAGGCCGGGTAAAGACAGAG | 53 |
| CTCF-2-2 | GGCGTCAGTCAAGTGATGG | TGTGAGGCGATTTAAACGTG | 53 |
| CTCF-3-1 | TGTTCATTTTTCAAAGTGAGCTTTA | ATGTGTGCGTCTGTTTCTGC | 53 |
| CTCF-9-1 | GCCATCTAGTGGTGCTGTTG | GTGAGCAATGTTCCCACCTT | 55 |
| CTCF-9-2 | GCCACCAGATGGCACTATTT | ACCGAATTGCCTCAGAACAG | 55 |
| CTCF-CHRX | TGCAGGAAATTCATGAGCTG | CATCCCTGGAACACAGATCC | 53 |
| FAM50B | GTGGTTCTCGTGGAGGTCAG | GCACCAATTTCCAGCATTTT | 52 |
| GRB10 | CGAAAGCCCTCCATGTCTAC | CCTTCCTGGTTCTTGCTCTG | 55 |
| IGF2 | CCTGCCTAGAGCTCCCTCTT | TTTCATATTCCGTGCCATGA | 52 |
| L3MBTL | GTTTCTGCCACCGTCCTG | CTGCGGTTTACGAGGCTTAG | 55 |
| MEG3-1 | CCAACGTCCCACGTCTCTAT | AGGGCTGAAAAGAAGGGAGA | 55 |
| MEG3-2 | GACTTTGGAGGCGTGATGTT | CCTGCTACAGGGAAAAGCAG | 55 |
| MEST | GTCCAGCACCGAACCTTTC | TTAAAAGGCATCCCTTACGC | 54 |
| NAP1L5 | TTAGAGGAGAAGCCGCAGAG | GCCCAGAAGATGTCAGGGTA | 55 |
| NDN | AAGGTGGAGTGCTTCTTCCA | GAGTTTGCGCTGGTCAAAG | 55 |
| NNAT | TTCTGGAAGGGGGCTAAGAT | TGGTCGAGAAAGAGGGTTTG | 56 |
| PEG10-1 | AGTCCTCGCGTGGTGAGTAT | GTTTTGGGGTACAGCCAGTG | 54 |
| PEG10-2 | CGCCTCCTCTTAGGTAGCC | CTGAAGGGAGGGGCTGAG | 54 |
| PLAGL1 | GGAGGACCTAAGCTGGGTGT | ATCGGGACAGCTGATAATGC | 55 |
| RB1 | GCGAGCAGCTCTCACCTCT | CCACGCTTCTACTCCCTGAA | 55 |
| RB-L1-1 | CAGGCCAGTGTGTGTGCG | AAAGCGCAATATTCGGGTGG | 55 |
| RB-L1-2 | GAGCTACGGGAGGACATTCA | CCTGAATCTGAACGTTGGCC | 55 |
| RB-L1-3 | CACGCAGCTGGAGATCTG | AATACCCTGCCGTGTGAGG | 55 |
| RB-L1-4 | TGCCTACAAGAGAAAGCAGGA | AGTCTTGCTAGCGGTCTATCA | 55 |
| RB-L1-5 | TGAAAACCGGCACAAGACAG | GCTGAGACGATGGGGTTTTC | 55 |
| RB-L1-8 | TGAAAACCGGCACAAGACAG | GCTGAGACGATGGGGTTTTC | 55 |
| SNRPN | CTTGACAATCCCCGAACACT | CACCCAGGGATGACTGACA | 55 |
| ZIM2 | ATATGCCACCAACCAACCAG | CCGAGTAGGCGCTGTTCTA | 55 |
| GOM-TDG | AAGTACAAACACCAGTGACAGA | AGCATCACCTCCCATAACCT | 53 |
| GOM-DNMT1 | CCTGATGCCCCTAGAACTGT | TATCTCGCCAACCTGACCTC | 59 |
| GOM-DNMT3A | ACTATAGACCAGGCGTGCTC | GCGGGTTGTGAGAAGGAATG | 59 |
| GOM-DNMT3B | TGTGTGCCCATAATGTTGCA | CCATTGTAGCCTCACCCTCA | 59 |
| LOM-TDG | CAAAGAGCTGTGATCATGCCA | GGTCATCCACTGCCCATTAG | 59 |
| LOM-DNMT1 | GACCGTGGACAGCTGACT | TGAGACTGAGCCTGAATCCA | 59 |
| LOM-TET3 | CATTGTGGGGTGTTGGTGG | CGCCCACCTCAGAAAACAG | 59 |
| LOM-MBD4 | TTATGCTGAAAACTGGCCTTG | AAATCAGTTAACGTTTCTCCACC | 59 |
| LOM-CPG-1 | TAGCTGAGCGGGAAGAAGC | GAGAACGACGGGGAGCTG | 60 |
| GOM-CPG-2 | GGTCTCCGTTCCCTTCTCC | TTCGGAAGTGGAAGAAGCCT | 60 |
| LOM-CPG-3 | GCTGCCTTCGAGCCTCTTC | CTAGGTGAAGAACGGGCAAC | 60 |
